# Using evolution as a tool: Replacing *corolla* in *Drosophila melanogaster* with its *Drosophila mauritiana* ortholog creates a novel hypomorphic allele

**DOI:** 10.1101/2025.06.11.659179

**Authors:** Stefanie Williams, Grace McKown, Zulin Yu, Cynthia Staber, Matt Gibson, R. Scott Hawley

## Abstract

In *Drosophila melanogaster* females, as in most organisms, the segregation of meiotic chromosomes depends on the proper distribution of crossovers along paired maternal and paternal chromosomes. In most cases, crossovers require the synaptonemal complex (SC), a conserved multi-protein structure that forms between homologous chromosomes in meiotic prophase I. Recent studies leveraging hypomorphic alleles suggest that the SC plays a more direct role in the distribution of crossover events. However, identifying additional hypomorphic mutations that avoid catastrophic phenotypes by partially disrupting the SC has been challenging. Here, to create a new hypomorphic allele of the *D. melanogaster* SC gene *corolla*, we used CRISPR/Cas9 to replace it with the coding sequence of its *Drosophila mauritiana* ortholog, yielding *corolla^mau^*. Since the amino acid sequence of SC proteins is rapidly diverging while maintaining the general tripartite structure of the SC, we hypothesized that this replacement would enable the assembly of the SC but show defects in crossover distribution. Indeed, at 25 °C *corolla^mau^* homozygous females exhibited full-length SC with defects in SC maintenance and crossover formation, resulting in moderate levels of chromosome missegregation. At 18 °C, SC maintenance was rescued, and recombination rates were improved, although they remained significantly lower than observed in wild type. Importantly, these phenotypes are less severe than observed in *corolla* null mutant flies, suggesting *corolla^mau^* is a hypomorphic allele. Unexpectedly, in homozygotes we also observed unique polycomplexes composed of the SC proteins Corolla and Corona but lacking the transverse filament protein C(3)G. Overall, we report a novel hypomorphic allele of *corolla* that will enable future studies on the role of the SC in crossover distribution. Further, the unique polycomplexes found in mutant flies may provide new insights into SC protein-protein interactions and SC architecture.

**Author Summary:** In many species, the success of sexual reproduction relies on a protein structure called the synaptonemal complex (SC). The SC forms between the maternal and paternal copies of chromosomes and functions to ensure crossing over. Most prior studies have used SC mutants that have grave defects, preventing the study of nuances in SC function. Here, we replace one of the SC genes in *Drosophila melanogaster* with the ortholog of a close relative, creating a new allele that displays a partial loss-of-function phenotype. At the standard rearing temperature, flies homozygous for this allele exhibit SC maintenance defects, a reduced number of crossover events, and aberrant chromosome segregation. In flies reared at a lower temperature, SC maintenance is rescued but the defects in recombination and chromosome segregation persist. We also found a unique SC protein aggregate in these flies. Altogether, this new mutant reflects a novel approach to study the structure and function of the SC.

## Introduction

Sexual reproduction relies heavily on the successful completion of meiosis. This specialized form of cell division reduces the chromosome number by half, ensuring that when an egg and sperm fuse, the correct ploidy is achieved. Mistakes during meiosis can lead to the improper segregation of chromosomes resulting in aneuploidy which is the leading cause of miscarriages and congenital syndromes such as trisomy 21 (1–3).

In most organisms, the formation of meiotic crossovers between homologous chromosomes is essential to ensure their proper segregation (4). These crossovers mature into chiasmata which hold the homologs together until they can be separated in anaphase I. Importantly, crossovers are not randomly distributed across the chromosomes. Instead, several mechanisms ensure their proper positioning (5). Crossover assurance ensures that a minimum of one crossover is established between each homologous chromosome pair (reviewed in (6)). In most species, the centromere effect suppresses the formation of crossovers within the centromere as well as the euchromatin in proximity to it. Precisely how proximal to the centromere a crossover can form differs between organisms (5,7–9). Similarly, crossovers are suppressed within and close to the telomeres, the ends of chromosomes generally consisting of repetitive non-coding DNA sequences and heterochromatin in most organisms (5,10). In addition, crossover interference enforces that two crossovers cannot occur in close proximity to one another, although the strength of this effect varies dramatically between species (5,7,11). The lengths of chromosome axes also play into the patterning of crossovers (5). As a result, crossover patterning is species-specific with observable differences even between closely related species. This is seen in *Drosophila melanogaster* and its close relatives *Drosophila mauritiana* and *Drosophila simulans,* both of which present with a larger map length and more centromere-proximal crossovers compared to *D. melanogaster* (12,13).

The precise role of the synaptonemal complex (SC) in the regulation of crossover formation and distribution remains poorly understood. The SC is a multi-protein structure that consists of two lateral elements connected to the homologous chromosomes and the central region with a central element that synapses them (Fig 1A). The protein components that make up the SC differ among established meiosis research organisms. To our current knowledge, the central region of the SC in *D. melanogaster* is comprised of three proteins: C(3)G, Corolla, and Corona (Cona) (Fig 1A) (14–16). C(3)G is the transverse filament protein (14). The C-termini of C(3)G are thought to interact with both lateral elements while the N-termini are thought to interact with the N-termini of another C(3)G dimer (14,17,18). The localization of Corolla within the SC has been described as transverse filament-like (15,18). Immunofluorescent labeling in combination with expansion microscopy and structured illumination imaging revealed that Corolla does not span the entire width of the SC from one lateral element to the other, but the signal is too wide for it to be considered a central element protein (18). Cona is known as the central element protein of *D. melanogaster* with a distinct localization in the very center of the SC (16,18).

**Fig 1.**
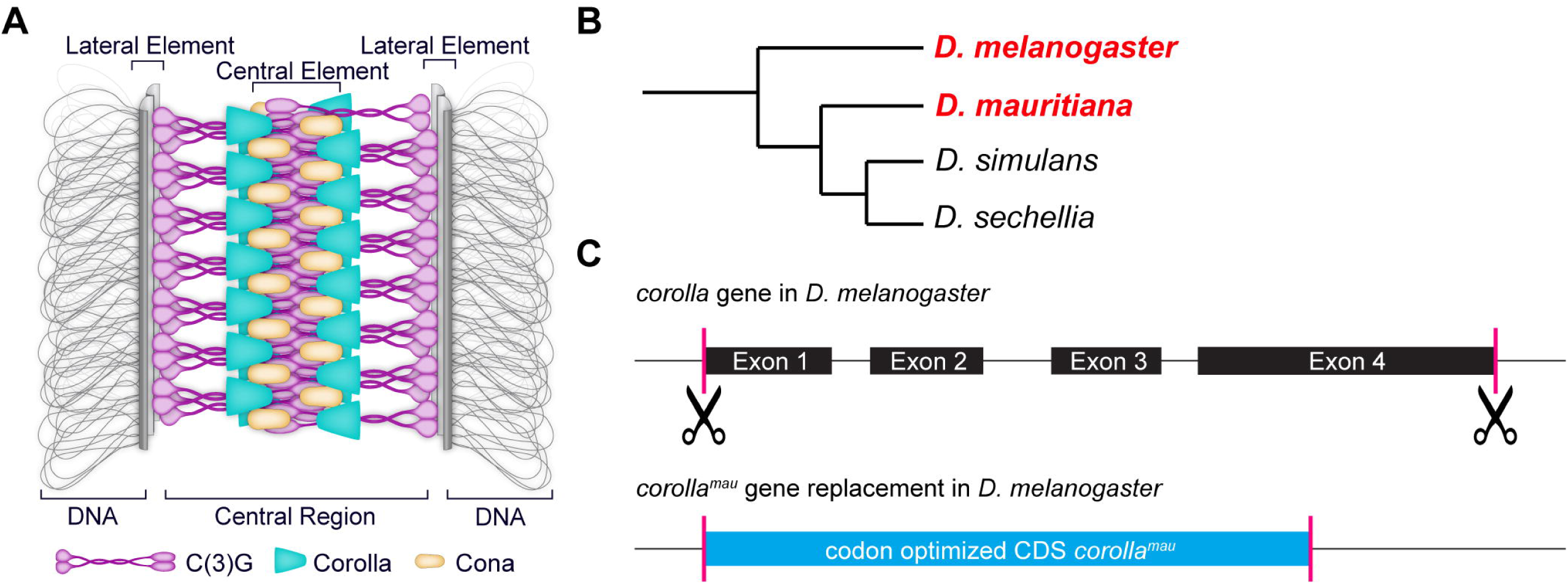
The replacement of the SC central region protein Corolla with its ortholog from *D. mauritiana*. (A) Model of the SC in *D. melanogaster*. The lateral element is connected to chromatin of the homologous chromosomes. Within the central region of *D. melanogaster* is the transverse filament protein C(3)G, Corolla, and the central element protein Cona. (B) The cladogram shows the closest relatives to *D. melanogaster* that are *D. mauritiana, D. simulans*, and *D. sechellia.* For the gene replacement of *corolla*, we chose *D. mauritiana*. (C) Schematic of how the gene replacement was performed. The gene *corolla* was excised from the start of exon 1 to the end of exon 4 using CRISPR/Cas9 technology. The codon optimized coding sequence (CDS) from *D. mauritiana* was then inserted in this same locus.

In most organisms, the fully assembled synaptonemal complex (SC) is required for meiotic crossover formation and thus accurate chromosome segregation. However, several exceptions have been found. In *Saccharomyces cerevisiae*, mutants missing the central element components Ecm11, Gcm2, or the majority of the transverse filament protein Zip1 N-terminus have higher levels of recombination despite lacking full-length SC (19). In *Arabidopsis thaliana*, double *null mutants* of the transverse filament proteins *zyp1a* and *zyp1b* resulted in asynapsis while crossover formation remained possible (20,21). The proper regulation of these crossovers was disrupted, resulting in the loss of crossover assurance for each chromosome, loss of crossover interference, and loss of typically observed sex differences in crossover rates (20,21). In *Oryza sativa*, the *zep1-3* mutant (expressing a truncated version of the transverse filament protein) shows elevated recombination rates and changes in crossover distribution (22). While these findings show that synapsis of the homologs is not a requirement for crossovers in some species, the observed changes in crossover numbers and their distribution do suggest a role of the SC in crossover patterning.

Interestingly, while the basic architecture of the SC is conserved across eukaryotes, the protein components that make up the SC are rapidly evolving (23–26). This rapid evolution of SC components presents a significant challenge in studying the role the SC plays in crossover formation as it makes the identification of conserved amino acids difficult. Conserved residues within a protein’s amino acid sequence are generally assumed to serve important functions, making them appropriate targets for mutagenesis (27). The rapid evolution suggests that changes to single amino acids can be tolerated without causing major meiotic defects and therefore do not cause easily recognizable phenotypes which can be recovered in genetic screens. A suppressor screen performed by Kursel et al. highlights this, as many of the recovered suppressors of the s*yp-1^K42E^* mutant phenotype had acquired mutations in *syp-1* and *syp-3,* but displayed no additional phenotypes (28). This indicates that many point mutations within SC genes are tolerated. As a result, most SC mutants recovered so far have been null alleles. While these mutants have been instrumental in understanding the essential role of the SC in crossover formation and chromosome segregation, the reliance on loss-of-function mutations comes with significant limitations. In most organisms, null alleles typically result in a complete failure of SC assembly, leading to an almost complete lack of crossover events between homologous chromosomes. This makes it challenging to genetically dissect the SC’s role in regulating the formation and distribution of crossover events.

Despite these challenges, there are key examples of partial loss-of-function alleles that have permitted the study of the SC’s role in recombination. In one case, targeted deletions in the *D. melanogaster* SC protein C(3)G suggested that the autosomes and the *X* chromosome require full-length SC at different times for their proper recombination (29). In *C. elegans,* lowering the expression levels of the transverse filament protein SYP-1 by RNAi resulted in a decrease of crossover interference (30). Additionally, two recent studies showed that mutations in the *C. elegans* SYP-4 C-terminus disrupt normal crossover patterning (19,20).

In an effort to create a novel hypomorphic allele by leveraging the rapid evolution of SC proteins, here we used CRISPR/Cas9 to replace *D. melanogaster corolla* with the ortholog from its close relative *D. mauritiana*. The two *Drosophila* species differ in their crossover landscape (12,13), and their respective Corolla proteins share 86.7% amino acid sequence identity. We hypothesized that the replacement allele, referred to as *corolla^mau^*, would result in a partial loss-of-function, as amino acid substitutions are found throughout the protein sequence. Indeed, we found that *corolla^mau^*females exhibit temperature-sensitive synapsis defects and global crossover defects. At a rearing temperature of 25 °C, *corolla^mau^* homozygous females assembled the SCs in early pachytene but showed early fragmentation and early disassembly. In addition, *corolla^mau^*mutants exhibited unusual polycomplexes (PCs) that contained Corolla and Cona but lacked the transverse filament protein C(3)G. These PCs suggest that the interaction of Corolla from *D. mauritiana* with Cona from *D. melanogaster* is functional but the interaction with C(3)G may be disrupted. We also observed significantly reduced levels of recombination on the *X* chromosome and, while not as severely affected, improper recombination levels and distribution of crossovers on the *2^nd^* chromosome. At 18 °C, SC maintenance was not negatively affected, correlating with an improvement of *X* chromosome recombination, although the levels were still significantly lower than in wild type (= *corolla^+^*). As recombination overall was decreased but not abolished, chromosome missegregation was significantly increased but lower than observed for the *corolla^null^*mutant. From this we infer that *corolla^mau^* is a hypomorphic allele as homozygotes present with proper SC assembly and maintenance at 18 °C, but crossovers and chromosome segregation are significantly impacted. Together, these results suggest that by replacing the Corolla protein from *D. melanogaster* with its *D. mauritiana* ortholog we successfully altered proper crossover formation. In addition, these data support the hypothesis that Corolla is not only a structural protein of the SC but plays an additional role in crossover formation. This is further supported by the fact that we find the C-terminus of Corolla to be enriched with the amino acid phenylalanine and the occurrence of LC3-interacting region (LIR) motifs, something that has been recently established as an important factor for the regulation of crossover formation (31).

## Results

### Replacing the *corolla* gene in *D. melanogaster* with its ortholog from *D. mauritiana* to create a hypomorphic allele

To circumvent the problem of identifying conserved amino acid motifs that we could target for the creation of a hypomorphic allele, we chose to replace the *D. melanogaster* gene encoding the central region protein, Corolla, with that of a close relative. The closest relatives of *D. melanogaster* are *D. mauritiana, D. simulans,* and *D. sechellia* (Fig 1B). Previous studies of recombination in *D. mauritiana* and *D. simulans* revealed that the total map length of *D. mauritiana* was 1.36 (13) to 1.8 (12) times more compared to *D. melanogaster*, and the total map length of *D. simulans* was 1.3 times more compared to *D. melanogaster* (12). Both species exhibit more recombination in the regions proximal to the centromere than *D. melanogaster* (12,13), but the strongest effect was seen in *D. mauritiana*. Therefore, we chose *D. mauritiana* as the donor for the replacement of the endogenous *corolla* gene in *D. melanogaster*, as it would allow us more easily to understand whether Corolla is directly involved in these differences of crossover patterning.

The amino acid sequence of Corolla from *D. mauritiana* shares 86.7% sequence identity with *D. melanogaster* Corolla (S1 Fig and S1 File). In total, there are 74 amino acid mismatches including a two amino acid gap (S1 Fig and S1 File). These mismatches have a similarity of 4.7%, meaning that a substituted amino acid is of similar chemical type. The DeepCoil2 program (33–35) indicates that Corolla from *D. mauritiana* has an overall higher probability for coiled-coil formation within the first 200 amino acids of the protein than *D. melanogaster* (S2A-B Fig). To form coiled-coils, a heptad repeat of seven amino acids must be present, usually denoted as a-b-c-d-e-f-g (36). The positions a and d within that heptad repeat are typically hydrophobic amino acids while positions e and g are typically charged or polar amino acids (36–38). The higher probability of coiled-coil formation in Corolla from *D. mauritiana* is likely because DeepCoil2 finds more a and d residues within the first 200 amino acids (S2A-B Fig). Importantly, the length of the region with coiled-coil probability is predicted to be of the same length within the first 200 amino acids of the proteins, and the arrangement of individually predicted coiled-coil domains is largely similar in spacing (S2A-B Fig). This is reminiscent of the findings by Kursel et al. which showed that within nematodes the length and location of coiled-coil domains is highly conserved as well as the overall protein lengths of orthologous SC proteins (26). Indeed, the overall length of the Corolla protein from both *Drosophila* species is strikingly similar with 554 amino acids in *D. melanogaster* and 556 amino acids in *D. mauritiana* (S1, S2 Figs and S1 File).

In addition to a higher probability of coiled-coil formation, we also find that Corolla from *D. mauritiana* has a slightly larger intrinsically disordered region (IDR), as well as a small additional IDR within the C-terminus of the protein according to an InterProScan (S2C-D Fig) (39). Specifically, the IDR in Corolla from *D. melanogaster* spans amino acids 308 – 432 while the first IDR in Corolla from *D. mauritiana* spans amino acids 305 – 462, and the second small IDR spans amino acids 476 – 495 (S2C-D Fig). IDRs have been established as important factors in the function of SC proteins. For example, a recent study showed that the IDR of the *C. elegans* central region protein SYP-4 is important for the regulation of crossovers and maintenance of biophysical characteristics of the SC (31,32). Importantly, both IDRs and coiled-coil domains are implicated in promoting phase-phase separation through multivalent interactions (26,40).

In light of the differences in protein architecture described above, we hypothesized that replacing *D. melanogaster* Corolla with its ortholog from *D. mauritiana* would allow for the structural integrity of the SC but could impact recombination. To generate *D. melanogaster* flies expressing the *D. mauritiana* Corolla protein, we used CRISPR/Cas9 to excise *corolla* exons 1-4 and inserted the codon-optimized coding sequence (CDS) of *D. mauritiana corolla* (Fig 1C). We wanted to insert the CDS in the endogenous *corolla* locus so that regulatory elements of *corolla* gene expression could act on the gene expression of the replacement. We also chose not to insert the introns from *D. mauritiana* to avoid problems caused by foreign intron sequences. The resulting allele, *corolla^mau^*, was then subjected to detailed phenotypic analysis.

In summary, we replaced the *corolla* locus in *D. melanogaster* with the codon-optimized CDS of its ortholog in *D. mauritiana.* The Corolla proteins from *D. melanogaster* and *D. mauritiana* are of similar length and the first 200 amino acids exhibit probability for coiled-coil formation. Importantly, the protein from *D. mauritiana* has an overall higher probability for coiled-coil formation in that region and additionally has a larger IDR in the C-terminus.

### *corolla^mau^* females exhibit early SC disassembly

Upon completion of the gene replacement, we first asked how this affected SC assembly, maintenance and disassembly. Within the germarium of *D. melanogaster*, meiotic progression can be followed spatially (Fig 2A). Using structured illumination microscopy (SIM) on *corolla^mau^* mutant germaria maintained at 25 °C, we observed that the SC assembled but was not maintained beyond early pachytene (Fig 2B). Specifically, compared with wild-type controls (=*corolla^+^*) we observed early SC disassembly in 1/35 nuclei in early pachytene, 14/16 nuclei in early-mid pachytene, and 5/5 of nuclei in mid pachytene (S3 Fig). To test whether the SC width was affected in *corolla^mau^* mutants we turned to stimulated emission depletion (STED) microscopy, which offers higher resolution than SIM. We used a C(3)G antibody that recognizes a region in the C-terminal globular domain of C(3)G (17). As the C-terminal region of C(3)G is localized in proximity to the lateral elements (17,18), the fluorescence signal should clearly resolve into two tracks. The two tracks of C(3)G in both *corolla^+^* and *corolla^mau^* mutants were clearly visible in the STED images (Fig 2D). We found that the SC width in *corolla^mau^*germaria was not significantly different (124 ± 16 nm) compared to the *corolla^+^*SC (124 ± 9 nm) as determined with a t test (p-value = 0.95) (Fig 2C). STED microscopy did reveal that only 14% of SCs in early pachytene were full-length while 64% of SCs already showed mild fragmentation (Figs 2D and S4), and 21 % of SCs show high levels of fragmentation (S4 Fig). By early-mid pachytene, 70% of SCs were mildly fragmented while the remaining 30% were highly fragmented (S4 Fig). The fact that fragmentation was apparent with STED but not SIM is likely due to the above-mentioned differences in the resolution of these microscopy techniques. Combined, these results show that that the replacement of Corolla from *D. melanogaster* with its ortholog from *D. mauritiana* allowed for the assembly of the SC, but that it could not be maintained beyond early pachytene, when SC fragmentation was already apparent.

**Fig 2.**
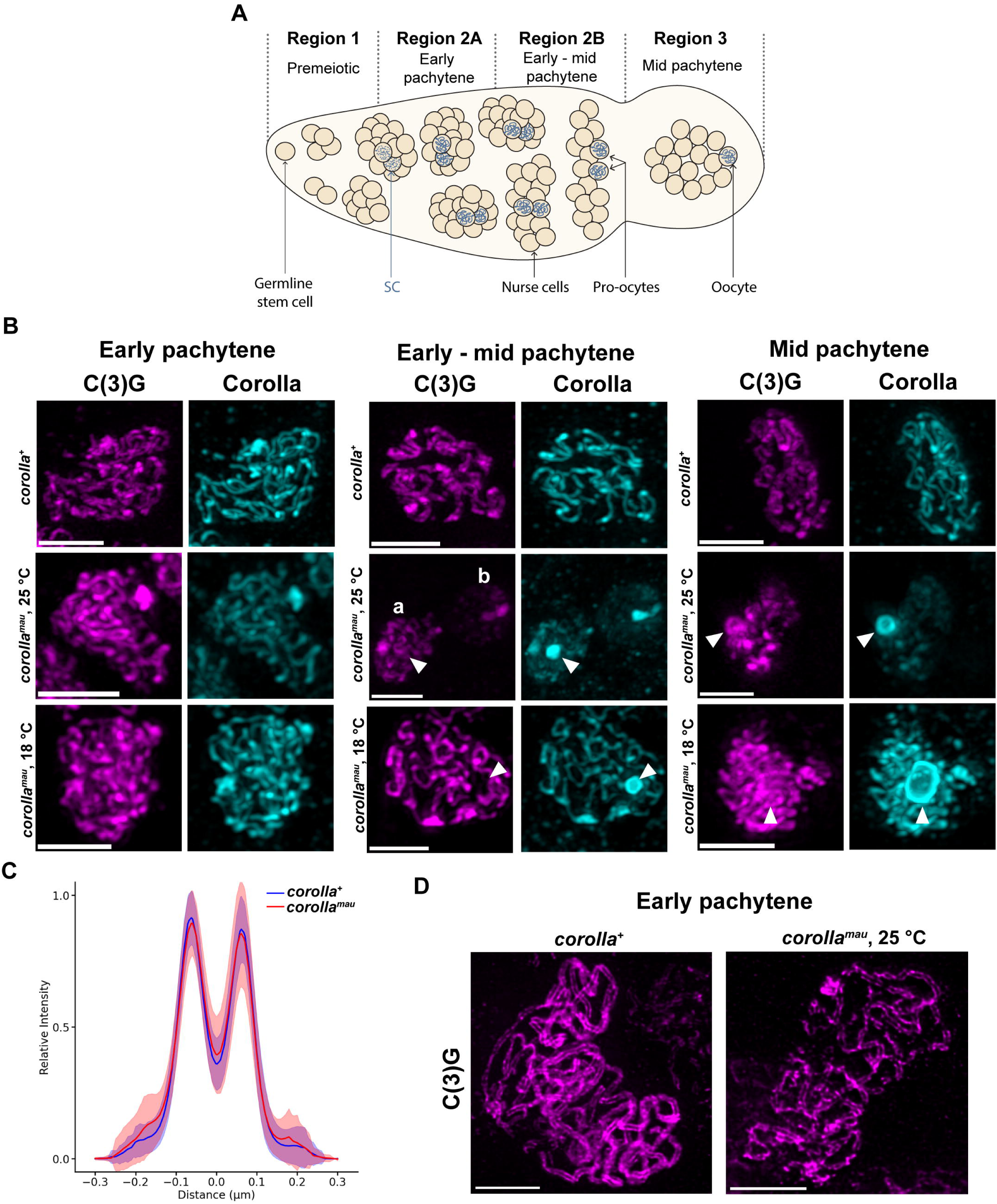
The *corolla^mau^* gene replacement causes SC maintenance defects. (A) Schematic of the *D. melanogaster* germarium. Germline stem cells are found in the premeiotic region 1. These proliferate to a 16-cell cyst at region 2A where two nuclei enter meiosis and assemble the SC in early pachytene and the remaining nuclei become nurse cells. As these cysts migrate along the germarium, they enter early-mid pachytene in region 2B where we still see two meiotic pro-oocyte nuclei. At mid pachytene, region 3, only one meiotic cell remains, the oocyte. The full-length SC is clearly visible from early pachytene to mid pachytene. (B). Immunofluorescence staining of the *corolla^+^* and *corolla^mau^* SC. In early pachytene, SC threads of both C(3)G (magenta) and Corolla (cyan) are present in *corolla^+^* and *corolla^mau^*. In early-mid pachytene, SC threads of C(3)G and Corolla are visible for *corolla^+^* and *corolla^mau^* reared at 18 °C. At the same stage, in *corolla^mau^* reared at 25 °C the SC is visibly disassembling (a) or has already disassembled (b). In mid pachytene, SC threads are visible in *corolla^+^* and *corolla^mau^* reared at 18 °C, but the SC in *corolla^mau^* flies reared at 25 °C is in the process of disassembling. Polycomplexes are visible in *corolla^mau^* nuclei regardless of rearing temperature (marked by arrowheads). Scale bars represent 2 µm length. (C) Relative fluorescence intensity of the two C-terminal C(3)G tracks over distance in *corolla^+^* and *corolla^mau^* reared at 25 °C. The SC is not significantly different in width compared between *corolla^+^* and *corolla^mau^*. (D) Deconvoluted maximum projection of the SC in *corolla^+^* and *corolla^mau^* imaged with STED microscopy. The two tracks of the C(3)G C-terminal antibody staining are visible. In *corolla^mau^*, SC mild fragmentation is visible. Scale bars represent 2 µm length.

### SC-dependent centromere clustering in *corolla^mau^* mutants

In *D. melanogaster,* the SC at the centromeres is different from the chromosome arms. Electron microscopy studies revealed that the central element has a less definite morphology and the lateral elements are surrounded by highly condensed chromatin making it more difficult to distinguish them from each other (41). In addition, at the beginning of prophase I the centromeres cluster into one or two masses (42). This process is called centromere clustering and requires the presence of C(3)G, Corolla, and Cona (15,42). Due to the observed early disassembly and SC fragmentation, we questioned whether centromere clustering would be affected in *corolla^mau^* flies. To analyze this, we stained germaria with an antibody against CID (Centromere Identifier, the *Drosophila* equivalent of CENP-A) and scored the number of clusters per nucleus (S5 Fig). In early pachytene, the average number of CID clusters was 1.9 ± 0.8 for *corolla^+^* and 1.8 ± 0.7 for *corolla^mau^* (S5 Fig). In early-mid pachytene, the average number of CID clusters was 1.9 ± 0.6 for *corolla^+^*and 2.0 ± 0.8 for *corolla^mau^* flies (S5 Fig). Lastly, in mid pachytene, the average number of CID clusters was 1.8 ± 0.5 for *corolla^+^*and 2.0 ± 0.7 for *corolla^mau^* females (S5 Fig). P values determined with an unpaired t-test indicated no statistical significance in any of these cases (S5 Fig). Therefore, we can conclude that SC-dependent centromere clustering is not affected in *corolla^mau^* mutants (S5 Fig). This shows that while Corolla from *D. mauritiana* causes defects with SC maintenance, other functions of Corolla like centromere clustering can be successfully completed.

### Polycomplexes in *corolla^mau^* flies do not consist of all SC protein components

In addition to the assembly of the tripartite SC that is not properly maintained, we noted the occurrence of polycomplexes (PC) in *corolla^mau^*flies (Fig 2B). PCs are structures consisting of SC proteins that assemble into repetitive structures that resemble the fine ultrastructure of the canonical SC (reviewed in (43)). While these are most often seen in mutant backgrounds that perturb the SC, well-studied examples of PCs include those induced by increased temperature in wild-type organisms (44,45). The spontaneous occurrence of PCs under standard conditions has also been reported, but it is not a common phenomenon (17,43). In *corolla^mau^* mutant germaria, 2/35 pro-oocytes at early pachytene, 11/18 pro-oocytes at early-mid pachytene, and 5/5 oocytes at mid pachytene contained PCs, which were not observed in *corolla^+^* (Figs 2B and S6A). Importantly, we found that 50% of PCs in *corolla^mau^* flies lacked the SC component C(3)G (Figs 2B and S6A-B). To our knowledge, PCs in flies usually consist of all SC central region proteins (43), which would make this the first observation of PCs lacking a central region component. The PCs are circular and appear to be hollow (Figs 2B and S6B-C). With this shape, the PCs are reminiscent of those seen in the c(3)*g^Cdel^* mutants reported by Jeffress et al. (46), as well as *sina^A4^/sina^Df^* mutants reported by Hughes et al. (47). However, all previously reported PCs contain C(3)G. To verify that the PCs were localized within the nucleus, we co-stained for the nuclear envelope protein Lamin and found that they were indeed nuclear (S6C Fig).

To summarize, we found that the replacement of *D. melanogaster* Corolla with its ortholog from *D. mauritiana* results in abnormal PC formation. Some of these PCs do not contain the transverse filament protein C(3)G, something that has never been observed before in flies. PCs may form due to the early disassembly of the SC, whereas the PCs lacking C(3)G may indicate a faulty interaction between C(3)G from *D. melanogaster* and Corolla from *D. mauritiana*.

### The *corolla^mau^* SC maintenance defect is temperature-sensitive

Due to the occurrence of PCs, and especially those without the transverse filament protein C(3)G, we wondered whether the observed SC disassembly was caused by an unstable interaction between *D. mauritiana* Corolla and the native *D. melanogaster* proteins Cona and C(3)G. Corolla and Cona have been previously shown to interact with each other in a yeast two-hybrid assay (15). Importantly, it is still unclear how these two proteins interact with C(3)G.

Previous studies have linked the formation of PCs with elevated temperature in several plant species and *C. elegans* (44,48). As the PCs observed in *corolla^mau^* mutants always contained Corolla and Cona, we infer that their interaction was not disrupted. While 25 °C is the standard rearing temperature for both *D. melanogaster* and *D. mauritiana*, we hypothesized that the 74 amino acid changes in *D. mauritiana* Corolla could destabilize its interaction with *D. melanogaster* C(3)G and Cona in a temperature-dependent manner. Hydrophobic multivalent interactions have been shown to be important for phase-phase separation (26,40,45), and a lowering of temperature leads to a reduction of molecular motion and the stabilization of condensates (49). Therefore, we asked if rearing *corolla^mau^* mutants at 18°C could rescue the early disassembly phenotype and the occurrence of PCs. Indeed, early SC disassembly was not observed in *corolla^mau^* flies reared at 18°C (Figs 2B and S7). Notably, PC formation was still observed in these germaria. Specifically, we observed PCs in 6/30 pro-oocytes at early pachytene, 8/10 at early-mid pachytene, and 3/3 at mid pachytene (S6A Fig). Interestingly, while the ratio of PCs with and without C(3)G was 50:50 (9/18 PCs for each type) for *corolla^mau^* reared at 25 °C, for those reared at 18 °C we observe that only 2/17 PCs contain C(3)G while 15/17 PCs did not (S6A Fig). From this we conclude that the formation of PCs is not caused by early SC disassembly in *corolla^mau^* mutants reared at 25 °C. We further observed fewer C(3)G-positive PCs in *corolla^mau^* females reared at 18 °C. This may suggest that Corolla from *D. mauritiana* and Cona from *D. melanogaster* are prone to form PCs with each other, but as the SC is not disassembling early at 18 °C, C(3)G is not readily available to join the PCs.

### Chromosome segregation and recombination are significantly affected in *corolla^mau^*females

Given the effects on SC assembly, we next tested how the *corolla* gene replacement affected chromosome segregation and recombination. In *corolla^mau^* flies, *X* chromosome missegregation increased significantly (Figs 3A and S8A). This effect was slightly less pronounced in flies reared at 18 °C (8.5% vs 10.8% at 25°C), but the difference was not statistically significant (Figs 3A and S8A). Importantly, at either temperature the rate of *X* chromosome missegregation was drastically lower than the 44.5% reported in *corolla* null mutant flies (15). *4^th^* chromosome missegregation was significantly lower in *corolla^+^* flies reared at 18 °C compared to 25 °C but, surprisingly, *corolla^mau^* females did not show significant differences compared to *corolla^+^* females (Figs 3A and S8A). This suggests that the proper segregation of the *4^th^* chromosome in control females was improved at 18 °C compared to 25 °C. We would therefore have expected to see an increase in *4^th^*chromosome missegregation in conjunction with the increase in *X* chromosome missegregation in *corolla^mau^* mutants. The small and mostly heterochromatic *4^th^* chromosome in *D. melanogaster* is, with very few exceptions, achiasmate and segregates via the distributive system (reviewed in (50,51)). The distributive system also ensures the proper segregation of any additional achiasmate chromosomes (51). Typically, with an increase in achiasmate chromosomes (e.g. through a mutant SC allele), *4^th^* chromosome missegregation is reported at half to two thirds level of *X* chromosome missegregation (15,47,52). This arises because the distributive system gets overloaded with achiasmate chromosomes and fails to segregate the *4^th^* chromosome. In *corolla^null^*flies for example, *X* chromosome missegregation was 44.5% and *4^th^*chromosome missegregation was 30.0% (15). It therefore remains unclear why *4^th^* chromosome missegregation is not affected in *corolla^mau^* flies.

**Fig 3.**
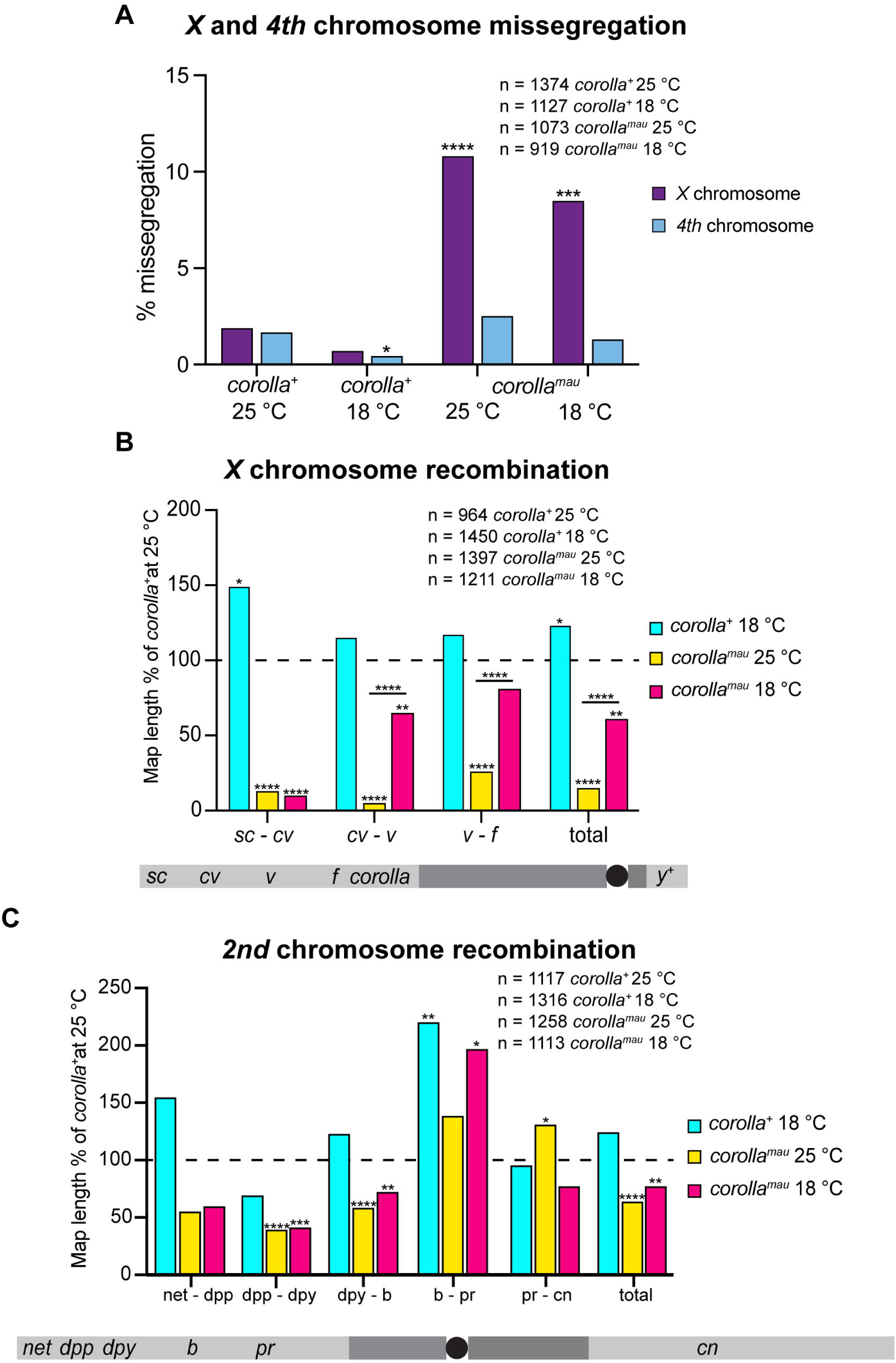
*corolla^mau^* females have modest defects in chromosome segregation and recombination. (A) *X* and *4^th^* chromosome missegregation. The percentage of missegregation is plotted for *corolla^+^* and *corolla^mau^* kept at both temperatures. (B) *X* chromosome recombination. The map length is plotted as a percentage of *corolla^+^* at 25 °C for each scored interval, as well as the total scored interval. A schematic of the *X* chromosome with the used marker genes scored as well as the location of *corolla* and *y^+^* is illustrated below the plot. (C) *2^nd^* chromosome recombination. The map length is plotted as a percentage of *corolla^+^* at 25 °C for each scored interval, as well as the total scored interval. Numbers of scored individual females for each experiment are provided for each genotype and temperature. A schematic of the *2nd* chromosome with the marker genes scored is illustrated below the plot. Statistical significance was calculated by Fisher’s exact test. Asterisks on top of bars indicate statistical significance in comparison to *corolla^+^* at 25 °C. Asterisks on top of horizontal lines indicate statistical significance comparing *corolla^mau^* at 25 °C to *corolla^mau^* at 18 °C. * means p ≤ 0.05, ** means p ≤ 0.01, *** means p ≤ 0.001, **** means p ≤ 0.0001.

Due to the elevated *X* chromosome missegregation in *corolla^mau^*flies, we next hypothesized that recombination would be significantly reduced due to the gene replacement. Considering that the rate of *X* chromosome missegregation in *corolla^mau^* mutants was not significantly different between the different rearing temperatures, we further hypothesized that recombination would be decreased to a similar extent in *corolla^mau^*flies reared at 18 °C and 25 °C. Indeed, *X* chromosome recombination was significantly affected by the gene replacement (Figs 3B and S8B). In *corolla^mau^*flies reared at 25 °C, all intervals scored for recombination across the *X* chromosome had dramatically reduced levels of recombination (Figs 3B and S8B). The reduction in crossing over ranged from 5% of the *corolla^+^* map length for the interval *cv-v* to 26% of the *corolla^+^* map length in the interval *v-f* (Figs 3B and S8B). In contrast, *corolla^mau^* flies reared at 18 °C had significantly higher levels of recombination in two intervals (65% of the *corolla^+^* map length in *cv*-*v* and 81% of the *corolla^+^* map length in *v-f*) than *corolla^mau^*flies raised at 25 °C (Figs 3B and S8B). In the interval *sc-cv*, however, recombination was still significantly lower than in *corolla^+^* with 10% of the *corolla^+^*levels (Figs 3B and S8B). Over the entire scoring interval on the *X* chromosome, *corolla^mau^*flies reared at both temperatures had significantly reduced levels of recombination. Specifically, *corolla^mau^* flies reared at 25 °C had only 15% of the *corolla^+^ X* chromosome map length and those reared at 18 °C only 61% (Figs 3B and S8B). Interestingly, *corolla^+^* flies reared at 18 °C had a significantly longer total map length (Figs 3B and S8B), suggesting that crossover interference is decreased at 18 °C compared to 25 °C. This aligns with previously published data showing that recombination in *D. melanogaster*, as well as *A. thaliana*, have a U-shaped response when plotting crossover rates over temperature (53,54). The results for *X* chromosome map lengths are also reflected in the exchange rank frequencies calculated with the Crossover Maximum Likelihood Estimation Tool (Table 1) (13). Exchange rank frequencies provide more accurate information about bivalents with no crossovers (E0), a single crossover (E1), two crossovers (E2), and three crossovers (E3) than the direct readout on crossover chromatid types from scoring crossovers (55). *X* chromosome E0 tetrads are highly elevated in *corolla^mau^* flies kept at both temperatures but values increased dramatically at 25 °C with 0.864 E0 frequency while the *corolla^+^* E0 frequency is 0.251 (Table 1). It is important to note that as the *corolla* locus is positioned between *cv* and the typical additional marker *y^+^* (located on the short arm), the interval *cv-y^+^* could not be scored, which is why the E0 frequency observed for *corolla^+^*may seem high compared to previously published data. While *X* chromosome E0 tetrads for *corolla^mau^* flies at 18 °C are still elevated (Table 1), they had significantly more recombination than those at 25 °C (Fig 3B and Table 1). The fact that *corolla^+^* flies reared at 18 °C had a significantly longer total map length is reflected in the E0 tetrads being less than for *corolla^+^* at 25 °C and both E1 and E2 tetrads are increased.

**Table 1:** Exchange rank frequencies on the *X* and *2^nd^* chromosomes.

| *X* chromosome | *corolla*<sup>+</sup>, 25 °C | *corolla*<sup>+</sup>, 18 °C | *corolla*<sup>mau</sup>, 25 °C | *corolla*<sup>mau</sup>, 18 °C |
| --- | --- | --- | --- | --- |
| E0 | 0.251 | 0.132 | 0.864 | 0.523 |
| E1 | 0.595 | 0.631 | 0.133 | 0.405 |
| E2 | 0.154 | 0.237 | 0.003 | 0.073 |
| 2<sup>nd</sup> chromosome | *corolla*<sup>+</sup>, 25 °C | *corolla*<sup>+</sup>, 18 °C | *corolla*<sup>mau</sup>, 25 °C | *corolla*<sup>mau</sup>, 18 °C |
| E0 | 0.224 | 0.074 | 0.515 | 0.447 |
| E1 | 0.691 | 0.817 | 0.434 | 0.449 |
| E2 | 0.078 | 0.073 | 0.032 | 0.090 |
| E3 | 0.007 | 0.036 | 0.019 | 0.014 |

Exchange rank frequencies for the *X* and 2<sup>nd</sup> chromosomes for *corolla*<sup>+</sup> and *corolla*<sup>mau</sup> females at 18 °C and 25 °C. Values were calculated with the Crossover Maximum Likelihood Estimation Tool (13).

Previously, it has been shown that distinct deletions in the *D. melanogaster* C(3)G coiled-coil region differentially impact recombination rates on the *X* chromosome and the autosomes (29). We therefore next examined chromosome *2* recombination rates in *corolla^mau^*mutants. A key difference observed between crossing over on the *X a*nd the *2^nd^*chromosome is that there were no significant differences between *corolla^mau^* flies reared at the two different temperatures in any of the scored intervals (Figs 3C and S8C). However, there were distinct differences between *corolla^mau^* and *corolla^+^* flies in specific scoring intervals. Specifically, in two intervals (*dpp*-*dpy* and *dpy*-*b*), *corolla^mau^*flies reared at either temperature had significantly reduced levels of recombination compared to *corolla^+^* reared at 25 °C (Figs 3C and S8C). For the interval *dpp-dpy, corolla^mau^* mutants reared at 25 °C had a map length that was 39% that of the observed map length in *corolla^+^*. In the same interval, the map length for *corolla^mau^* mutants reared at 18°C only improved non-significantly to 41% of the *corolla^+^* map length (Figs 3C and S8C). Scoring of the interval *dpy-b* of *corolla^mau^* females reared at 25 °C and 18 °C revealed 58% and 72%, respectively, of the *corolla^+^* map length. In the interval *b*-*pr*, *corolla^mau^* flies reared at 18 °C had significantly higher levels of recombination, as did *corolla^+^* flies at the same temperature (Figs 3C and S8C). In the interval *pr*-*cn*, *corolla^mau^* flies reared at 25 °C had significantly higher levels of recombination (Figs 3C and S8C). This is especially interesting because *pr-cn* spans the centromere of chromosome *2*, indicating that crossovers in the region around it have increased. However, we do not know how close these crossovers were to the pericentromeric heterochromatin. Across the entire scoring interval on the *2^nd^* chromosome, *corolla^mau^* flies reared at both temperatures had significantly reduced levels of recombination with 64% of the *corolla^+^* map length for those reared at 25 °C and 77% for those reared at 18 °C (Figs 3C and S8C). As a result, E0 tetrads for the *2^nd^* chromosome were elevated for *corolla^mau^* mutants at both temperatures, accompanied with a large decrease in E1 tetrads in comparison to *corolla^+^* flies (Table 1). Again, E0 tetrads for *corolla^+^* flies reared at 18 °C were dramatically reduced compared to those reared at 25 °C (0.074 vs 0.224), and both E1 and E3 tetrads were increased (Table 1).

In summary, *X* chromosome missegregation was significantly increased in *corolla^mau^* mutants compared to *corolla^+^*regardless of the temperature they were raised in, while *4^th^* chromosome missegregation was not significantly altered. On the *X* chromosome, a significant improvement in crossover rates was observed in *corolla^mau^* flies reared at 18 °C compared to those reared at 25 °C, but overall map lengths on the *X* chromosome and the *2^nd^* chromosome were significantly reduced in *corolla^mau^* mutants at both temperatures. These data suggest that the distributive system maintains its ability to segregate the *4^th^* chromosome but sporadically fails to segregate the *X* chromosomes.

### The C-terminus of Corolla shares sequence features with SYP-4 in *C. elegans* and SIX6OS1 in vertebrates

The fact that SC disassembly was rescued by rearing flies at 18 °C while the total map lengths on the *X* and *2^nd^* chromosomes remained reduced may suggest that Corolla plays an additional role important to recombination. Recent studies of SYP-4 in *C. elegans* (31,32) suggest it may be an ortholog of the known vertebrate SC protein SIX6OS1. The C-terminus of both proteins is rich in phenylalanine compared to the entire protein sequence, and both proteins feature LC3-interacting region (LIR) motifs. Neves et al. (31) and Köhler et al. (32) report that specific perturbations within the SYP-4 C-terminus significantly disrupt the regulation of crossover formation. We therefore investigated whether *Drosophila* Corolla is a SIX6OS1 ortholog and found that the C-terminus was indeed enriched with phenylalanine. In *D. melanogaster,* 10% of the last 89 amino acids of Corolla were phenylalanine, compared to 4% in the entire protein sequence. Similarly, in *D. mauritiana* Corolla 9% of the last 89 amino acids were phenylalanine compared to 4% in the entire amino acid sequence. Additionally, we found two LIR motifs in Corolla protein sequences from *D. melanogaster* and its *D. mauritiana* ortholog using the Eukaryotic Linear Motif (ELM) resource for functional sites in proteins (S9 Fig) (56). One of these motifs is shorter by two amino acids in *D. melanogaster*, which means that through the gene replacement, this motif was extended, and it additionally contains 3 amino acid substitutions. One of these is an amino acid substitution of a phenylalanine with an aspartic acid at the aligned residue 351 (residue 349 in the *D. melanogaster* protein sequence). Phenylalanines in LIR motifs were shown to have a special functional importance in SYP-4 (31). While it is currently unclear whether these changes to the LIR motif can explain the recombination defects seen in *corolla^mau^* mutants, future experiments will hopefully shed light on this.

In summary, we found that the Corolla C-terminus is enriched in phenylalanines and contains two LIR motifs, two factors that have recently established a connection between SYP-4 and SIX6OS1, and are important for the regulation of crossover formation (31,32). Together, this may suggest that Corolla, SYP-4 and SIX6OS1 are orthologs and that Corolla plays an additional role in crossover formation.

## Discussion

### The *corolla^mau^* gene replacement created a hypomorphic mutation

The SC assembles in early meiosis I to connect homologous chromosomes and ensure crossover formation. Most mutations available to research the SC are null alleles that prevent proper SC assembly and do not allow for direct studies on how the SC is involved in the process of crossover placement. By replacing the *corolla* gene in *D. melanogaster* with its ortholog from *D. mauritiana*, we have created the first partial-loss-of-function allele of *corolla*. When reared at 25 °C, these *corolla^mau^* flies presented with fragmentation and early disassembly of the SC. While this defect shows that the SC consisting of Corolla from *D. mauritiana* and C(3)G and Cona from *D. melanogaster* cannot be properly maintained, the SC-dependent process of centromere clustering was not affected. Furthermore, the early disassembly of the *corolla^mau^* females was rescued by lowering the rearing temperature to 18 °C, indicating that problems in SC maintenance did not arise from irreconcilable incompatibility between Corolla from *D. mauritiana* and C(3)G and Cona from *D. melanogaster.* In addition, we observed PCs in *corolla^mau^*reared at both temperatures, some of which contain Corolla and Cona but lack the transverse filament protein C(3)G. In *corolla^mau^* flies reared at 18 °C, we observed that 88% of PCs lack C(3)G, while this number is decreased to 50% when reared at 25 °C. *X* chromosome missegregation was significantly increased in *corolla^mau^* females to 9-11% at both rearing temperatures, but importantly, these missegregation levels are moderate compared to the *X* chromosome missegregation rate of 44.5% in *corolla^null^*females (15). Furthermore, *corolla^mau^*mutants showed no elevated rates of *4^th^*chromosome missegregation at any of the two rearing temperatures compared to *corolla^+^*, while *corolla^null^* mutants had a *4^th^* chromosome missegregation rate of 30.0% (15). In accordance with the increased rate of *X* chromosome missegregation, the map lengths of the *corolla^mau^ X* and *2^nd^*chromosomes were significantly reduced compared to *corolla^+^ X* and *2^nd^* chromosomes at both temperatures. *X* chromosome recombination levels were increased in flies reared at 18 °C relative to those at 25 °C but still significantly lower than in *corolla^+^* flies. It is important to emphasize that while overall map lengths were significantly reduced compared to *corolla^+^* flies, crossovers were not as depleted as in *corolla^null^* females (15). It is evident that the gene replacement in *corolla^mau^* flies caused significant recombination defects compared to *corolla^+^* flies but these defects were again not as severe as in the *corolla^null^*mutant. Therefore, by replacing the *corolla* allele in *D. melanogaster* with the *corolla* coding sequence from *D. mauritiana*, we established a hypomorphic allele. The partial-loss-of function becomes strikingly clear in *corolla^mau^* flies reared at 18 °C. While the SC maintenance defect was rescued at this lower rearing temperature, recombination was still significantly lower than *corolla^+^*females. This further suggests that the SC and/or the protein Corolla play a role in the regulation of crossover formation. One such role is further supported by the fact that Corolla shares conserved amino acid sequence features with the *C. elegans* SC protein SYP-4 and the vertebrate ortholog SIX6OS1. These features have been directly linked to crossover regulation in *C. elegans*.

### The SC in *corolla^mau^* flies is unstable at 25 °C

We have shown that the SC disassembles early in the *corolla^mau^*germaria at 25 °C, a phenotype that is rescued at 18 °C (Figs 2B and S3-5). These data suggest that the introduction of Corolla from *D. mauritiana* into the SC with Cona and C(3)G from *D. melanogaster* creates an instability that is intolerable at 25 °C but tolerable at 18 °C. Protein-protein interactions between Corolla and Cona have been shown in the past by the yeast two-hybrid assay (15), but it is still unclear how or if C(3)G directly interacts with these two proteins. Potentially, some of the 74 amino acid mismatches between Corolla from *D. melanogaster* and *D. mauritiana* are important for this interaction with C(3)G. The SC has been shown to have features of a biomolecular condensate (45) and lower temperatures are known to stabilize biomolecular condensates as hydrophobic interactions are strengthened through a reduction in molecular motion (49). This may explain why the early disassembly of the SC is rescued in *corolla^mau^* females at 18 °C. One potential explanation is that some of the 74 amino acids are important for the proper establishment of connections within the SC to maintain it. While at 25 °C mismatches introduced through the gene replacement are intolerable and lead to the disassembly of the SC, the interactions are generally stabilized at the lower temperature of 18 °C, allowing for its proper maintenance.

### Unique polycomplexes have been identified in *corolla^mau^* germaria

We report the first occurrence of PCs in flies that lack the transverse filament protein. Potentially, this tells us how the Corolla protein from *D. mauritiana* interacts with the other SC proteins in *D. melanogaster*. As described above, the coiled-coil domain of Corolla from *D. mauritiana* has an overall higher probability to form such coiled-coils and its IDR is longer compared to Corolla from *D. melanogaster* (S2 Fig). While the functional implications of both factors are currently unknown, one hypothesis is that these cause Corolla from *D. mauritiana* to form PCs. Future experiments will be needed to determine the functional impact of the higher probability for coiled-coil formation and longer IDR on PC formation.

Our initial hypothesis was that SC disassembly and the formation of PCs without C(3)G were both caused by an inability to interact with C(3)G. That hypothesis is clearly incorrect. While lowering the rearing temperature rescued SC disassembly, it did not affect total PC number. Instead, we observed an increase in the proportion of PCs without C(3)G. This may be the reason why we did not see C(3)G associating with more PCs in *corolla^mau^* females reared at 18°C. Potentially, as the SC is not disassembling, the majority of C(3)G proteins stay within the canonical SC and do not associate with the PCs, while a significant portion of Corolla and Cona proteins do associate in PCs, with or without C(3)G.

As an alternative explanation for the formation of PCs in *corolla^mau^* mutant germaria, we compared linear recognition motifs present on the Corolla proteins from *D. melanogaster* and *D. mauritiana* using the ELM resource for functional sites in proteins (S1 Table) (56). Important to note here is that the *corolla* gene theoretically encodes for two isoforms of Corolla with only a three amino acid difference (S10 Fig). Currently, it is unclear whether both isoforms are expressed, and functional studies should help clarify this. We also do not know whether the linear motifs mentioned below are functionally important for Corolla and the SC. Corolla isoforms A and B in *D. melanogaster* contain five and six functional sites, respectively, that are unique to the *D. melanogaster* protein. Candidates that could potentially create problems are the absence of a COP1 E3 ligase binding degron motif, a SUMO interaction site (both found on isoform A and B), as well as an APC/C destruction box motif (only on isoform B) on the Corolla protein from *D. mauritiana* (S1 Table). The lack of these motifs may affect transcriptional regulation and protein degradation. Importantly, the gene *sina* also codes for an E3 ligase. While the *sina* mutants studied in Hughes et al. are mutations in the gene body (47), the lack of the required motif would likely also result in functional problems. In the Corolla protein of *D. mauritiana* there are two unique motifs, one of which is an additional motif for the deubiquitinase Usp7. While it was shown that the knockdown of Usp7 has detrimental effects on SC maintenance, recombination, and fertility (57), it is unclear what effects an additional Usp7 motif would have. Future research is needed to elucidate whether any of these motifs have functional consequences and are the cause of the observed PCs in *corolla^mau^* females or the early SC disassembly when these flies are kept at 25 °C.

### Recombination and chromosome missegregation in *corolla^mau^* females indicate that Corolla plays a role in crossover formation

In *corolla^mau^* flies reared at 25 °C we see dramatically reduced levels of recombination on the *X* chromosome (Fig 3B) while *X* and *4^th^* chromosome missegregation is moderate in comparison to the *corolla^null^* mutants (15) (Fig 3A). Crossover levels on the *X* chromosome were increased in *corolla^mau^*females reared at 18 °C, coinciding with the rescue of SC maintenance defects at the standard rearing temperature. These results align with data from the deletions in the coiled-coil region of C(3)G, which suggested that full-length SC is needed from early to early-mid pachytene (29). As the SC in *corolla^mau^* germaria reared at 25 °C starts to show fragmentation in early pachytene and disassembles in early-mid pachytene, *X* chromosome recombination is decreased by a great extent.

Importantly, the results for recombination and missegregation seen in *corolla^mau^* flies are not identical with those reported for c(3)*G* mutants by Billmyre and Cahoon et al. (29). The early SC disassembly in two of the c(3)*G* partial deletions resulted in significant reductions of *X* chromosome recombination, which coincided with no significant increase in *X* or *4^th^*chromosome missegregation. This was likely because *X* chromosome crossovers failed to form due to the early disassembly of the SC, while autosomal crossovers were changed in a different manner. The SC disassembly in these partial c(3)*G* deletions leads to improper crossover distribution and specifically a significant increase in centromere proximal crossovers (29) which was not observed in *corolla^mau^*females. Directly comparing the values for *X* and *2^nd^* chromosome crossovers as well as *X* and *4^th^* chromosome missegregation between Billmyre and Cahoon et al. with results from *corolla^mau^*females, it becomes strikingly clear that Corolla may play a role in crossover formation in addition to its structural function (S11 Fig) (29). Even in the c(3)*G* partial deletion females with the lowest rates of *X* chromosome recombination (c(3)*G^CC3^*), *X* and *4^th^*chromosome missegregation were not as dramatically increased as in *corolla^mau^*females (S11 Fig).

This additional role for Corolla is further supported by our findings that Corolla shares amino acid sequence features with *C. elegans* SYP-4 and the vertebrate protein SIX6OS1 (31,32). Importantly, while one of the LIR motifs has distinct differences between *D. melanogaster* and *D. mauritiana*, the number of LIR motifs is the same in Corolla in both species. Therefore, it is unclear whether any of the other amino acid substitutions created by the gene replacement are the cause for crossover defects in *corolla^mau^* mutants or if it is these differences in the LIR motif. The similarities between Corolla, SYP-4, and SIX6OS1 make a striking case for their consideration as orthologous proteins.

To conclude, here we generated the first hypomorphic allele of *D. melanogaster corolla* by replacing it with the coding sequence of its *D. mauritiana* ortholog. The partial loss-of-function nature of this mutation becomes strikingly clear when we compare phenotypes observed in *corolla^mau^*flies to the *corolla^null^* mutant. Null mutations are integral to provide binary answers to questions like: “Is the SC required for crossover formations?” With novel alleles like *corolla^mau^*, we can now begin to untangle the complexities of the SC’s involvement in crossover formation and how the SC functions as a complex. This opens many avenues for future research. The failure to maintain the SC beyond early pachytene at the standard rearing temperature of 25 °C, which is rescued by rearing at 18 °C, opens the possibility that the SC in *corolla^mau^* females is destabilized due to amino acid mismatches. Future research could identify which of the 74 mismatches between Corolla from *D. melanogaster* and *D. mauritiana* contribute to this instability. The formation of PCs lacking C(3)G adds to this question but raises additional questions. When *corolla^mau^*flies are reared at 18 °C, why do we see a decrease in PCs that have C(3)G compared to 25 °C? Is C(3)G simply less readily available to join the PCs with the full-length SC maintained? Lastly, the global but moderate decrease of crossovers together with the moderate increase in *X* chromosome missegregation suggest that Corolla may play a direct role in crossover formation. Our ability to establish a connection between Corolla, SYP-4, and SIX6OS1 strengthens this possibility, but future work will be needed to address which of the differences between *D. melanogaster* and *D. mauritiana* Corolla result in crossover defects.

## Materials and Methods

### Stock maintenance

All flies used here were maintained on standard food. Flies for the experiments at 25 °C or 18°C were reared at their respective temperature.

### Construction of CRISPR stock

*w^1118^hyb5b-8, corolla^mauEx1^* was made by WellGenetics Inc. using a modified method of CRISPR/Cas9-mediated genome editing published by Kondo and Uedo in 2013 (58). The upstream CRISPR/Cas9 target site was selected to be +18 nt away from the start codon of *corolla* in *D. melanogaster* (CCCATAAATTCTTTGACAAG[TGG] with the PAM site in square brackets). The resulting upstream guide RNA primer oligos were the following: Sense oligo5’-CTTCGCCCATAAATTCTTTGACAAG and Antisense oligo 5’-AAACCTTGTCAAAGAATTTATGGGC. The upstream homology arm consisted of 1000 bp, specifically those -1000 nt to -1nt upstream of the *corolla* start codon in *D. melanogaster.* The downstream CRISPR/Cas9 target was selected to be -21 nt away from the stop codon of *corolla* in *D. melanogaster* (ATGTACGCCGAGGCGGCAGT[AGG] with the PAM site in square brackets). The resulting downstream guide RNA primer oligos were the following: Sense oligo5’-CTTCGATGTACGCCGAGGCGGCAGT and Antisense oligo5’-AAACACTGCCGCCTCGGCGTACATC. The downstream homology arm consisted of 993 bp, specifically those +4 nt to +996 nt downstream of the stop codon of *corolla* in *D. melanogaster.* The knock-in cassette consisted of the codon optimized coding sequence of *corolla* in *D. mauritiana* and a selection marker to facilitate screening. The selection marker consisted of Piggy Bac 5’ terminal repeats, the artificial 3xP3 promoter, DsRed2, SV40 3’UTR and Piggy Bac 3’ terminal repeats. For codon optimization of the *corolla* CDS the IDT DNA codon optimization tool ((https://www.idtdna.com/CodonOpt) with the gBlocks option was used. The codon optimization was necessary to ensure robust expression of the *D. mauritiana corolla* gene in *D. melanogaster* and to circumvent inefficiencies in the CRISPR/Cas9 mediated replacement of the *corolla* gene due to similarities in DNA sequence. The knock-in cassette and homology arms were cloned into the pUC57-Kan vector using sequence and ligation independent cloning and subsequently transformed using a standard protocol. The correct plasmid was selected using colony PCR and sequencing and subsequently used for microinjections into 226 wild-type embryos of the strain *w^1118^hyb5b-8*. 97 G0 males were outcrossed to females with the balancer *X* chromosome *FM7a*. Out of 72 viable crosses, 5 F1 lines were positive for the visible 3xP3 DsRed selection marker. F1 lines were tested for the insertion of the full knock-in cassette by obtaining genomic DNA from single flies and using PCR primers that should only provide an amplicon if the cassette is present. To test the insertion of the cassette in the 5’ region, the forward PCR primer is designed to anneal to a region outside of the upstream homology region (5’-GCATGTCCTTGGATGTCAGA) and the reverse primer is designed to anneal to a region within the *corolla* CDS from *D. mauritiana* (5’-GAACTGGGCCATCTGAGTGT). To test the insertion of the cassette in the 3’ region, the forward PCR primer is designed to anneal to the Piggy Bac 3’ terminal repeat (5’-TTTGACTCACGCGGTCGTTA) and the reverse primer is designed to anneal to a region outside of the downstream homology arm (5’-CTTCTGCTTGAGCAGCTCGT). Samples from two individuals presented with positive amplification of both PCR reactions and no amplification was found in the negative control *w^1118^hyb5b-8.* One of these samples was excised from the agarose gel and via sequencing the correct sequences upstream and downstream were confirmed. To test whether the backbone of the pUC57-Kan vector was inserted into the genome as well, two additional PCR reactions were performed. For the upstream region, the forward primer was designed to anneal to the region outside of the homology arm within the plasmid, i.e. the backbone of the plasmid (5’-CAACTGTTGGGAAGGGCGAT) and the reverse primer from the previous PCR reaction, annealing within the *corolla* CDS from *D. mauritiana*, was reused (5’-GAACTGGGCCATCTGAGTGT). For the downstream region, the forward primer annealing to the Piggy Bac 3’ terminal repeat (5’-TTTGACTCACGCGGTCGTTA) was reused, and the reverse primer was designed to anneal to the downstream region of the homology arm within the plasmid, i.e. the backbone of the plasmid (5’-CATTAGGCACCCCAGGCTTT). No amplification was found in either of the two individuals that were positive for the cassette insertion as well as the negative control *w^1118^hyb5b-8*. Amplification was found from the positive control, the donor plasmid. Positive excision of the selection marker using the Piggy Bac transposase was examined by another PCR reaction using a forward primer annealing to the 3’ region of the codon optimized *corolla* CDS from *D. mauritiana* (5’-GGAAGATGAATTCCCGATGA) and a reverse primer annealing to the downstream homology arm (5’-ACGGCACTTTCGAACTGAAT). All samples showed successful excision of the selection marker as the unexcised line had a larger amplicon and the negative control *w^1118^hyb5b-8* had no amplicon. Via sequencing it was confirmed that one TTAA motif was left in the 3’ end of the codon optimized *corolla* CDS from *D. mauritiana* resulting in a silent mutation. Two isogenized and balanced stocks were received from WellGenetics Inc. and the line *w^1118^hyb5b-8, corolla^mauEx1^* was used for crosses to establish the genotypes used in the different experiments.

### Chromosome missegregation assay

To assess *X* and *4^th^* chromosome missegregation, we crossed single virgin female flies of the genotypes y *w/ y w; sv^spa^-^pol^*(= *corolla^+^*) and y w *corolla^mauEx1^/* y w *corolla^mauEx1^; sv^spa^-^pol^* (= *corolla^mau^*) with males of the genotype *X^Y, ln(1)EN, v f B; C(4)RM, ci ey^R^.* For experiments at 25 °C, mating was set up for 5 days and progeny was subsequently scored for 18 days from the day of hatching. For experiments at 18 °C, mating was set up for 7 days and adults were moved to fresh vials that were kept at 18 °C for further mating while initial vials were moved to 25 °C for 7 days. Subsequently, progeny was scored for 8 days from the day of hatching. Calculations to determine the percentage of *X* and *4^th^*chromosome missegregation were performed as before (59,60). P values to determine statistical significance were calculated with the Fisher’s Exact test.

### Recombination assays

To assay recombination on the *X* chromosome we crossed single virgin females of the genotypes *y w/ y^1^ sc^1^ w^+^ cv^1^ v^1^ f^1^* (= *corolla^+^*) and *y w corolla^mauEx1^/ y^1^ sc^1^ w^+^ cv^1^ v^1^ f^1^ corolla^mauEx1^* (= *corolla^mau^*) with males of the genotype *y^1^ sc^1^ cv^1^ v^1^ f^1^ car^1^/ B^S^Y*. For experiments at 25 °C, mating was set up for 5 days and progeny was subsequently scored for eight days from the day of hatching. For experiments at 18 °C, mating was set up for 7 days, adults were moved to fresh vials that were kept at 18 °C for further mating while vials containing progeny were moved to 25°C for 7 days. Subsequently, only the female progeny were scored for 8 days from the day of hatching for the intervals *sc-cv, cv-v, v-f*.

To assay recombination on the *2^nd^* chromosome we crossed single virgin females of the genotypes *y w/ y; net^1^ dpp^ho^ dpy^ov1^ b^1^ pr^1^ cn^1^/ +* (= *corolla^+^*) and *y w corolla^mauEx1^/ w corolla^mauEx1^; net^1^ dpp^ho^ dpy^ov1^ b^1^ pr^1^ cn^1^/ + (= corolla^mau^)* to males of the genotype *w^+^/ Y; net^1^ dpp^ho^ dpy^ov1^ b^1^ pr^1^ cn^1^/ net^1^ dpp^ho^ dpy^ov1^ b^1^ pr^1^ cn^1^*. For experiments at 25 °C, mating was set up for 5 days and progeny was subsequently scored for eight days from the day of hatching. For experiments at 18 °C, mating was set up for 7 days and adults were moved to fresh vials that were kept at 18°C for further mating while vials containing progeny were moved to 25 °C for 7 days. Subsequently, only the female progeny were scored for 8 days from the day of hatching for the intervals *net-dpp, dpp-dpy, dpy-b, b-pr,* and *pr-cn*.

### Immunostaining of germaria

For microscopy, germaria of the genotypes *w/w* and *y w/ y w; pol* were used as *corolla^+^* and there were no differences observed between these. For *corolla^mau^* germaria of the genotypes *w corolla^mauEx1^*/ *w corolla^mauEx1^*and *y w corolla^mauEx1^*/ *y w corolla^mauEx1^*; pol where also no differences were observed. Germaria were immunostained as in Lake et al. (61), except that the samples were incubated with secondary antibodies for 2 hours instead of 4 hours and mounting was performed in ProLong Glass which was allowed to harden for at least 24 hours in the dark at room temperature. Primary antibodies used were affinity-purified rabbit anti-Corolla (1:2000) (15), mouse anti-C(3)G 1A8-G2 (17) (1:500), guinea pig anti-Cona (16) (1:500), mouse anti-lamin Dm0 ADL84.12 (1:100) (Developmental Studies Hybridoma Bank, Iowa), and rat anti-CID (1:500) (gift from the Sunkel lab (62)). Secondary antibodies were used at 1:500 and included Alexa Fluor 488 goat anti-mouse (ThermoFisher, A-11001), Alexa Fluor 555 goat anti-rabbit (ThermoFisher, A-21428), Alexa Fluor 647 goat anti-guinea pig (ThermoFisher, A-21450), Alexa Fluor 647 goat anti-mouse IgM (ThermoFisher, A-21238), and Alexa Fluor 594 goat anti-mouse (ThermoFisher, A-11032). DNA was stained with DAPI at a final concentration of 1ug/ml for 10 minutes.

### Imaging and Image Analysis

#### Structured Illumination Microscopy

Structured Illumination Microscopy (SIM) was performed with a 63x, N.A 1.40 Plan-Apo oil objective on a ZEISS Elyra 7 microscope. Alexa Fluor 647 signal was excited with 5% laser power for 100 ms, Alexa Fluor 555 signal was excited with 3% laser power for 80 ms, Alexa Fluor 488 signal was excited with 3% laser power for 50 ms, and DAPI signal (405 nm) was excited with 15% laser power for 100 ms. Images were taken with a z-stack size of 1.0 nm z-stack size and 15 phases. Images were then processed with the SIM tool in the Zeiss software. The parameter was set to “adjust” and the sharpness was set to 10.5 for the channels 647, 568, and 488 and set to 3 for the 405 channel. The method was set to “best fit”.

#### Stimulated Emission Depletion Imaging and SC width measurement

Stimulated Emission Depletion (STED) imaging was performed with a 100x, N.A. 1.40 oil objective on a Leica Sp8 Gated STED microscope. Alexa Fluor 594 signal was excited with a pulsed white light laser (80 MHz) tuned to 594 nm with 8% laser power. The pinhole was set to 0.6 Airy unit (AU) and nuclei were zoomed onto with a factor of 12.5. The z step size was set to 0.14 nm. The signal was depleted with a pulsed STED 775 nm laser at 90% power and the delay time between excitation and depletion was set to 600 ns. Images were acquired in 2D mode and averaged 8 times using the line average mode. Emission photons were detected with an internal Leica HyD hybrid detector with a time gate between 0.6 - 6 ns.

Raw images were deconvolved using the Huygens professional deconvolution software (version 22.04). The width of C(3)G was measured following the method described by Billmyre and Cahoon et al. (29). Well-defined SC regions that were roughly parallel to the xy plane were manually selected, and intensity profiles were obtained by averaging across a three-pixel-wide stripe oriented perpendicular to the SC. For C(3)G distance measurements, these profiles were fitted with double Gaussian functions, and the distance between the two peak positions was calculated.

#### Widefield imaging on DeltaVision microscopy system

Images with CID staining were acquired on an inverted DeltaVision microscopy system (GE Healthcare) with an Olympus 100× objective (UPlanSApo 100×, NA 1.40) and a high-resolution CCD camera or an Applied Precision OMX Blaze microscope equipped with a PCO Edge sCMOS camera. Images were deconvolved prior to analysis using SoftWoRx v. 7.2.1 (Applied Precision/GE Healthcare) software.

#### Figure preparation

For figure preparation of SIM and STED images, the region of interest was selected and duplicated. The duplicated image was z projected with the maximum projection setting. The brightness and contrast were minimally adjusted to ensure proper visualization of the SC and PCs. Channels were then split and lookup tables were applied to specific channels (see figure legends for figure-specific information on lookup tables). Scale bars of 2 µm were added to the C(3)G channel.

#### Scoring of CID foci, PCs, disassembly, and fragmentation

Scoring of CID foci was completed using ImageJ v.1.54f software with minimal brightness and contrast adjustment for ease of foci visualization. To score the occurrence of PCs and disassembly, SIM images were manually scored for each region (early pachytene, early-mid pachytene, mid pachytene) for the number of PCs relative to numbers pro-oocytes at the distinct stage as well as how many PCs contained or did not contain C(3)G. Disassembly was also scored with SIM images by counting pro-oocytes with visibly disassembling SC at distinct stages. Fragmentation was scored in maximum projected and deconvoluted STED images.

### Data availability

Original data underlying this manuscript can be accessed from the Stowers Original Data Repository at https://www.stowers.org/research/publications/libpb-2560

## Acknowledgments

We would like to thank Cathleen M. Lake and Stacie E. Hughes for fruitful discussions throughout the project. We would also like to thank Tolkappiyan Premkumar for the discussion that led to our discovery of temperature-sensitive phenotypes in *corolla^mau^*. A very big thank you to Cathleen M. Lake and Stacie E. Hughes for their help with the manuscript and additional thanks to Helen R. Horkan and Kimla J. Virden for feedback on the writing. Thanks also go to Angela Miller for the illustration of the synaptonemal complex. S. W. would also like to thank Amanda Kroesen and Cathy McKinney for assistance with the SIM and STED microscopes respectively. R.S.H was an American Cancer Society Researcher.

## Supporting information captions

S1 Fig. Pairwise alignment of the Corolla amino acid sequence in *D. melanogaster* (isoform A) and *D. mauritiana*. White background indicates 100% identity, black background indicates a mismatch.

S2 Fig. (A) DeepCoils2 result for Corolla from *D. melanogaster* showing the probability for coiled-coil formation on the y-axis and the amino acid position on the x-axis. At the top of the graph the a and d positions of amino acids within heptad repeats is indicated. (B) DeepCoils2 result for Corolla from *D. mauritiana* showing the probability for coiled-coil formation on the y-axis and the amino acid position on the x-axis. At the top of the graph the a and d positions of amino acids within heptad repeats is indicated. (C) InterProScan result of Corolla from *D. melanogaster* with the IDR highlighted. (D) InterProScan result of Corolla from *D. mauritiana* with the two IDRs highlighted.

S3 Fig. Quantification of full-length SC or disassembling SC in *corolla^+^*and *corolla^mau^* flies based on SIM images.

S4 Fig. Quantification of full-length SC, mild and high degrees of fragmentation in *corolla^+^* and *corolla^mau^* flies based on STED images.

S5 Fig. Quantification of the average number of CID foci per nucleus in early pachytene, early-mid pachytene, and mid pachytene.

S6 Fig. Polycomplexes in *corolla^mau^*. (A) Quantification of the occurrence of polycomplexes (PCs) from early pachytene to mid pachytene and of PCs with or without C(3)G for each genotype at both temperatures. (B) Same SIM images of *corolla^mau^* at 25 °C as shown in Fig 2B but including Cona from early to mid pachytene, highlighting co-localization of Corolla and Cona in a PC at early-mid pachytene but lacking C(3)G. Scale bars represent 2 µm length. (C) SIM images of *corolla^+^* , and *corolla^mau^* at both temperatures immunostained for C(3)G, Corolla, and Lamin.

S7 Fig. Quantification of disassembly in *corolla^+^* and *corolla^mau^* reared at 18 °C.

S8 Fig. (A) Raw numbers of total flies scored (n) for each genotype at each temperature and numbers of flies scored for *X* and *4^th^*chromosome missegregation. The percentage of missegregation scored is listed in brackets. (B) *X* chromosome recombination. Raw numbers of total flies scored (n) for each genotype at each temperature and numbers of recombinant flies recovered for each interval. Map length for each interval is provided in brackets. % of crossover chromatid types is provided for noncrossover chromatids (% NCO), single crossover chromatids (% SCO), and double crossovers (% DCO) recovered. (C) *2^nd^*chromosome recombination. Raw numbers of total flies scored (n) for each genotype at each temperature and numbers of recombinant flies recovered for each interval. Map length for each interval is provided in brackets. % of crossover chromatid types is provided for noncrossover chromatids (% NCO), single crossover chromatids (% SCO), double crossovers chromatids (% DCO), and triple crossover chromatids (% TCO) recovered.

S9 Fig. The two LIR motifs found in Corolla in both *D. melanogaster* and *D. mauritiana*. In the first LIR motif, the three amino acid mismatches are highlighted in black and a gap of two amino acids is indicated with horizontal lines.

S10 Fig. Pairwise alignment of isoform A and isoform B of Corolla in *D. melanogaster*. White background indicates 100% identity, black background indicates a mismatch.

S1 File. Pairwise sequence alignment showing that there are 74 mismatches, 86.7% sequence identity and 4.7% similarities between Corolla from *D. melanogaster* and *D. mauritiana*.

S1 Table. Functional sites found in Corolla isoform A and isoform B in *D. melanogaster* and Corolla in *D. mauritiana.* Sites marked in light blue are shared between *D. melanogaster* and *D. mauritiana.* Sites marked in pink are unique to *D. melanogaster*. Sites marked in yellow are unique to *D. mauritiana*.

## Notes

### Competing Interest Statement

The authors have declared no competing interest.

