## Supplementary figures and images for "Using evolution as a tool: Replacing *corolla* in *Drosophila melanogaster* with its *Drosophila mauritiana* ortholog creates a novel hypomorphic allele"

### Supplemental Figure 1

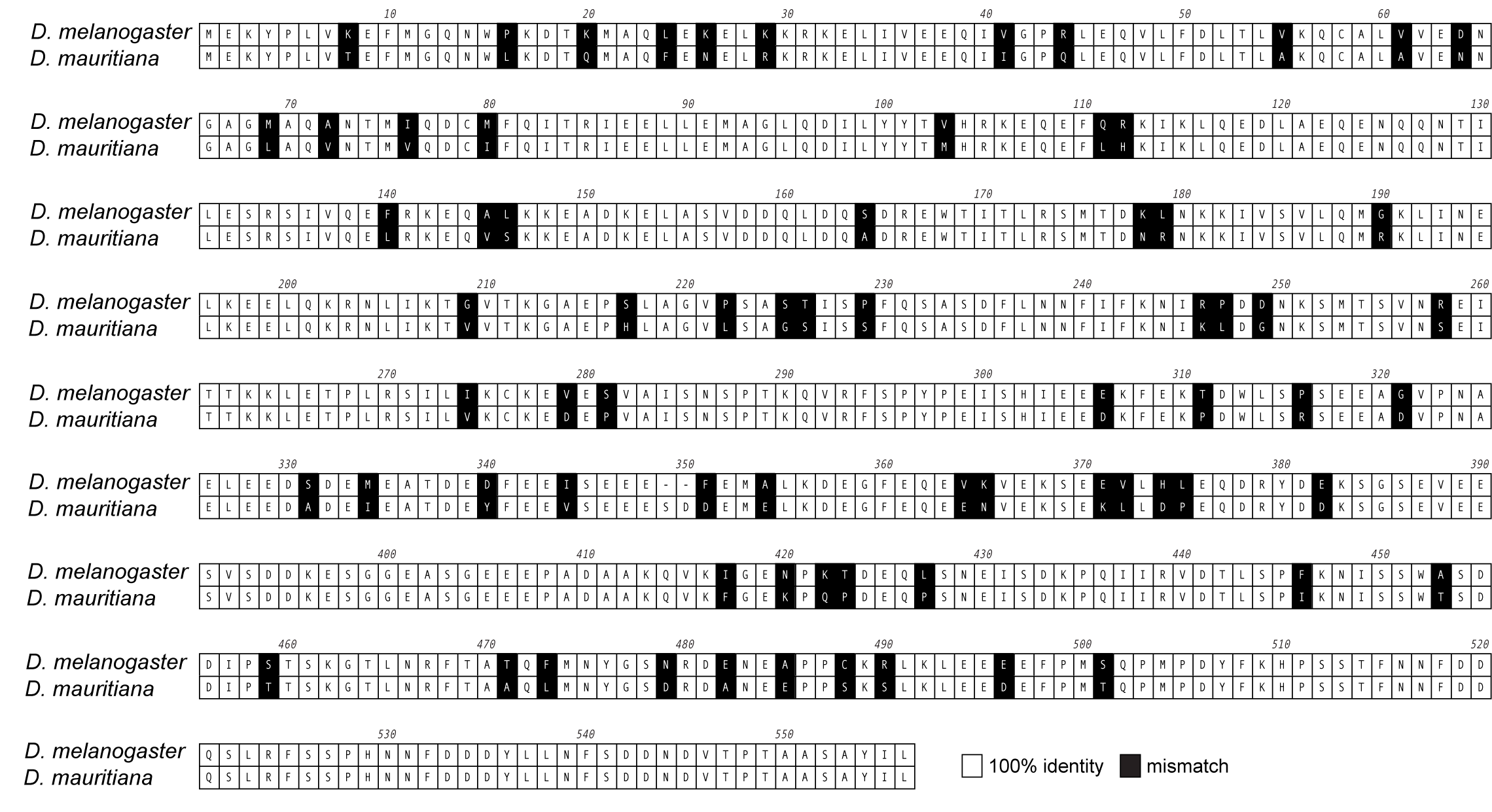

### Supplemental Figure 2

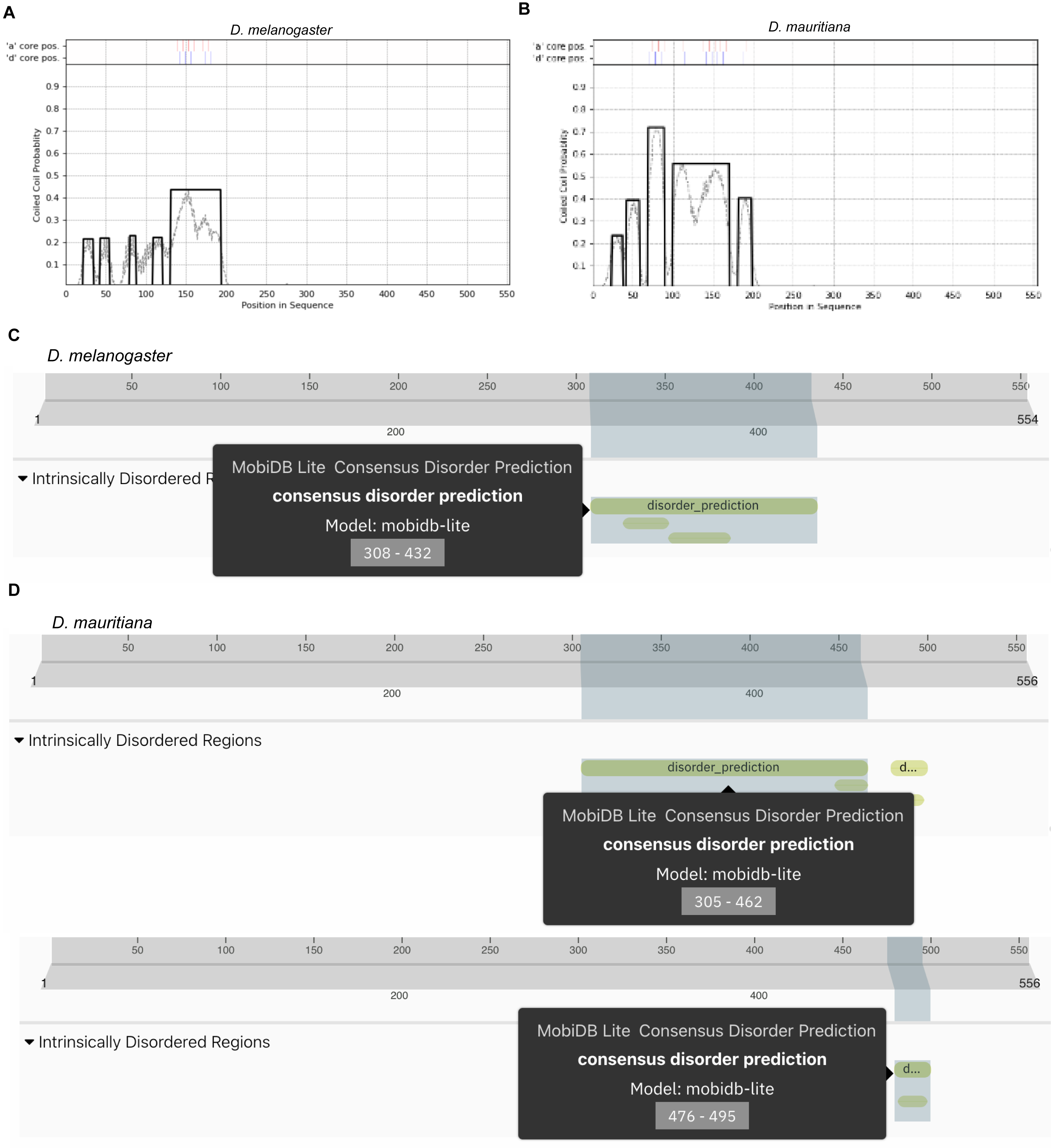

### Supplemental Figure 3

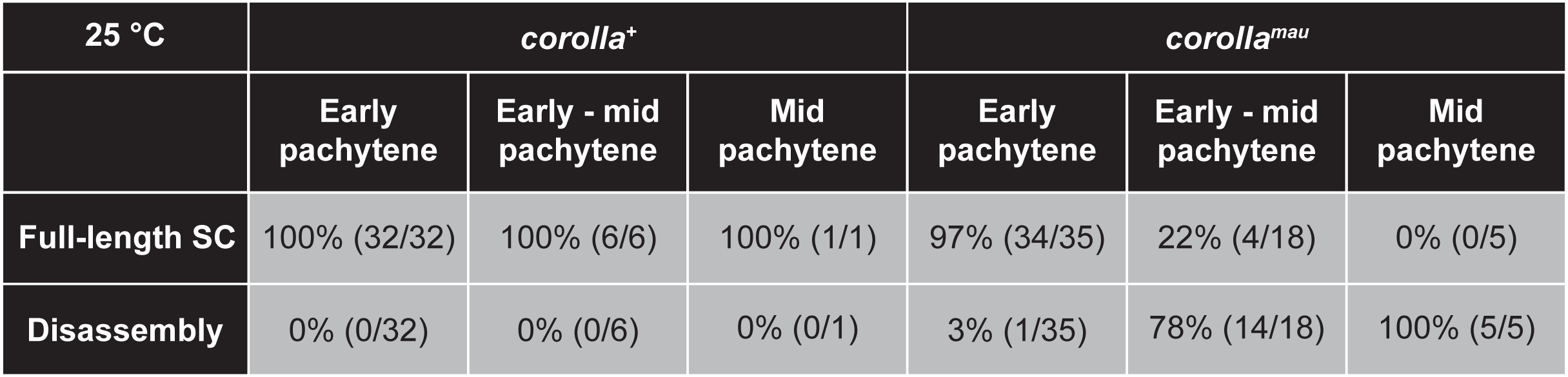

### Supplemental Figure 4

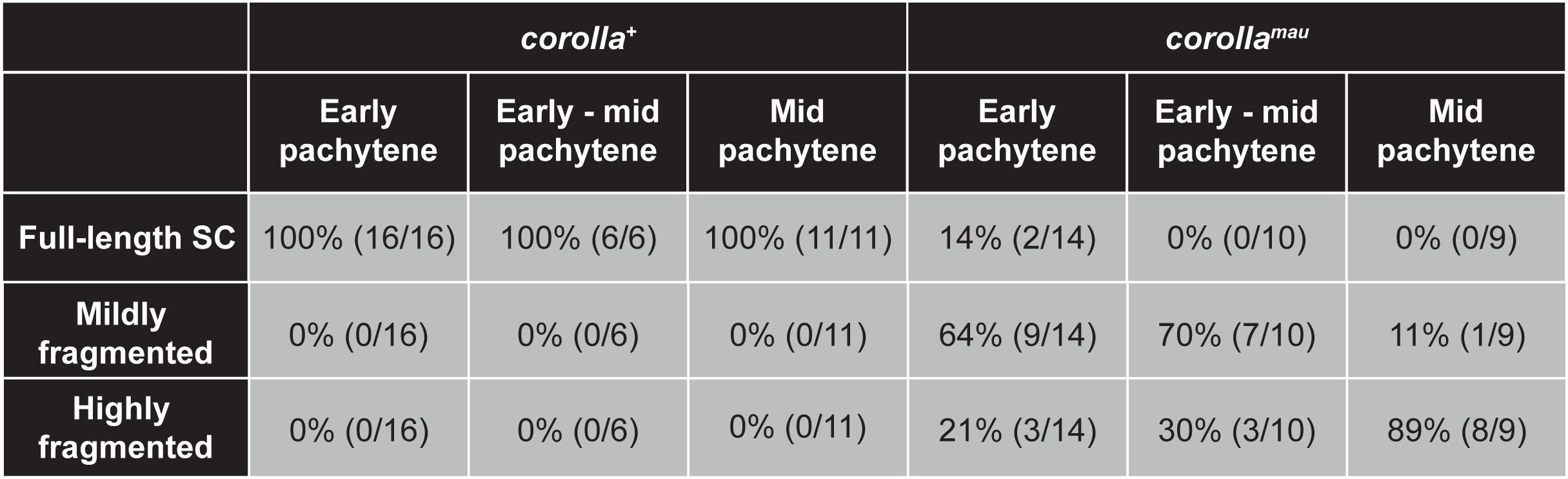

### Supplemental Figure 5

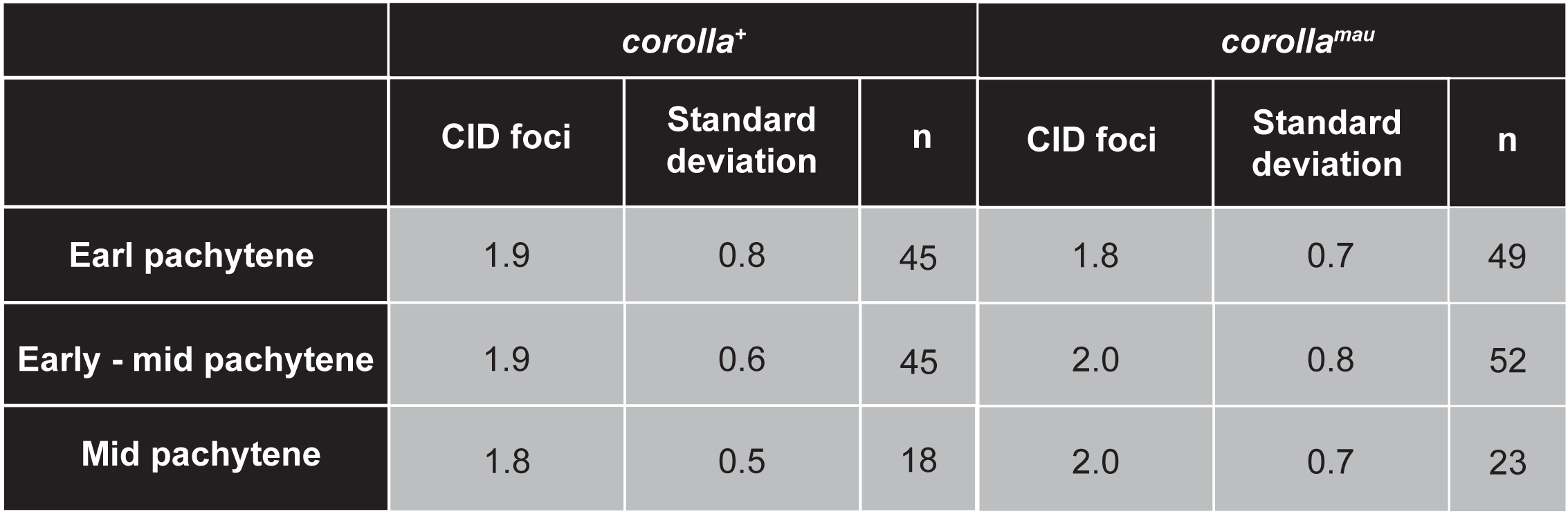

### Supplemental Figure 6

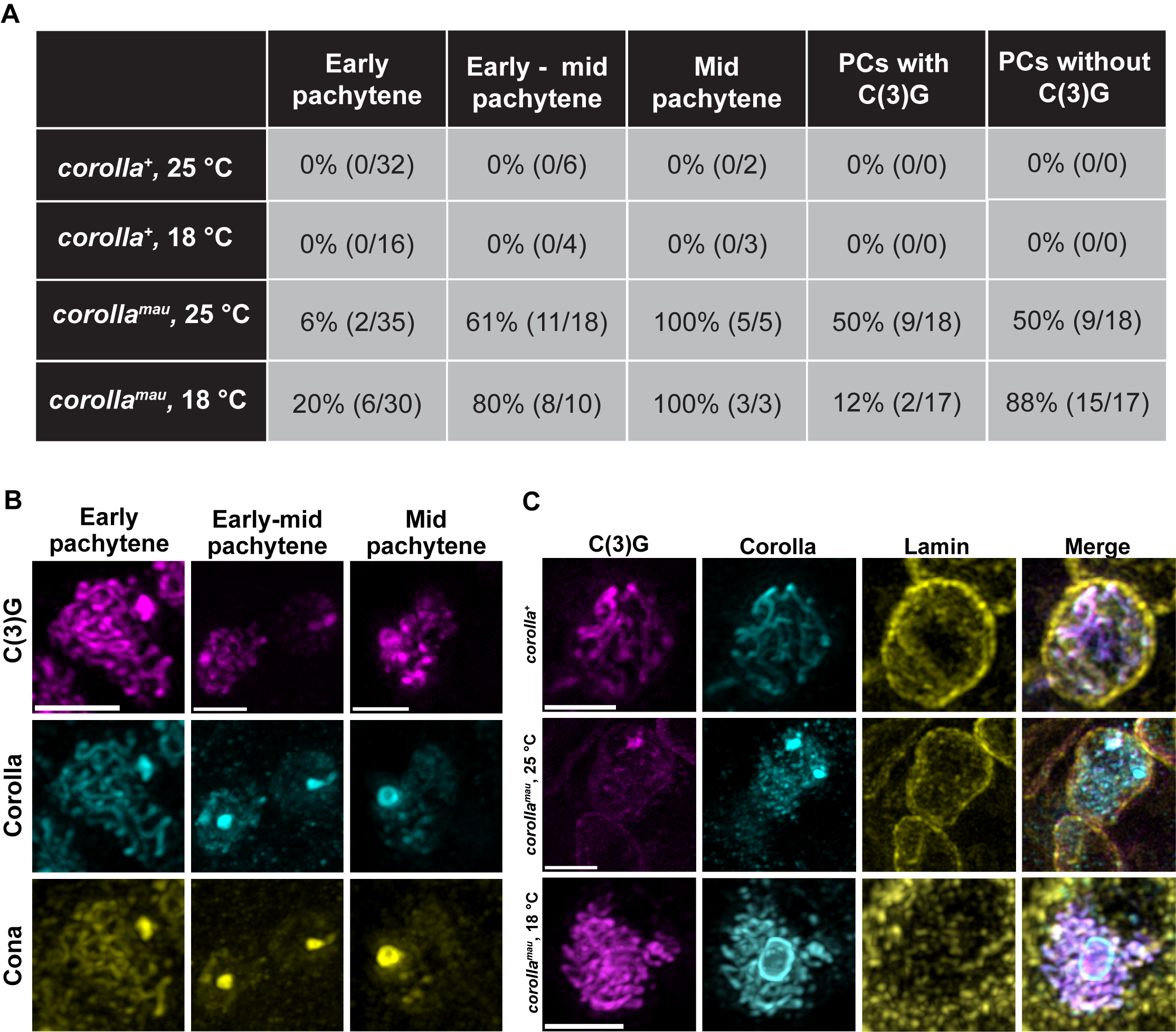

### Supplemental Figure 7

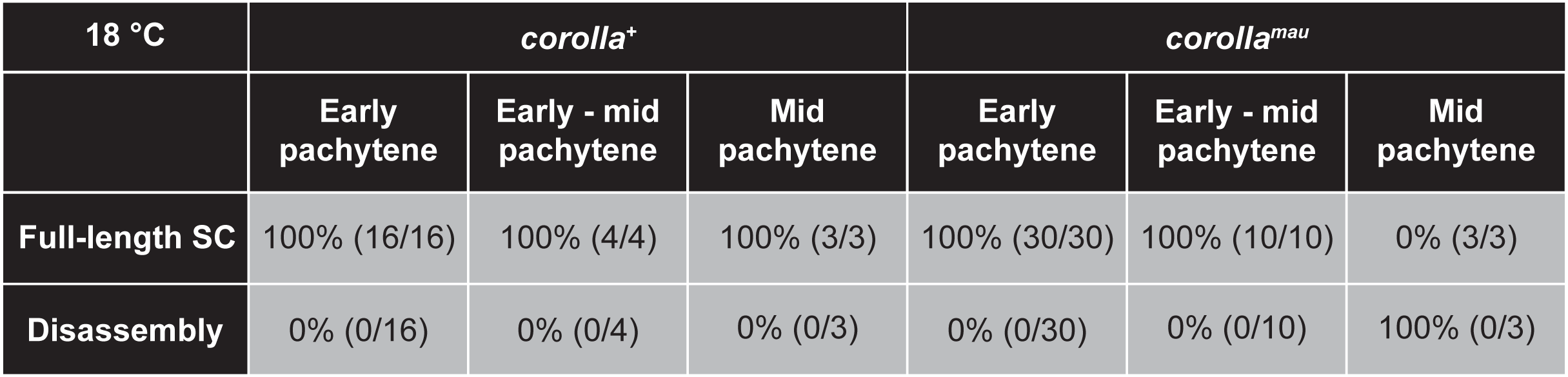

### Supplemental Figure 8

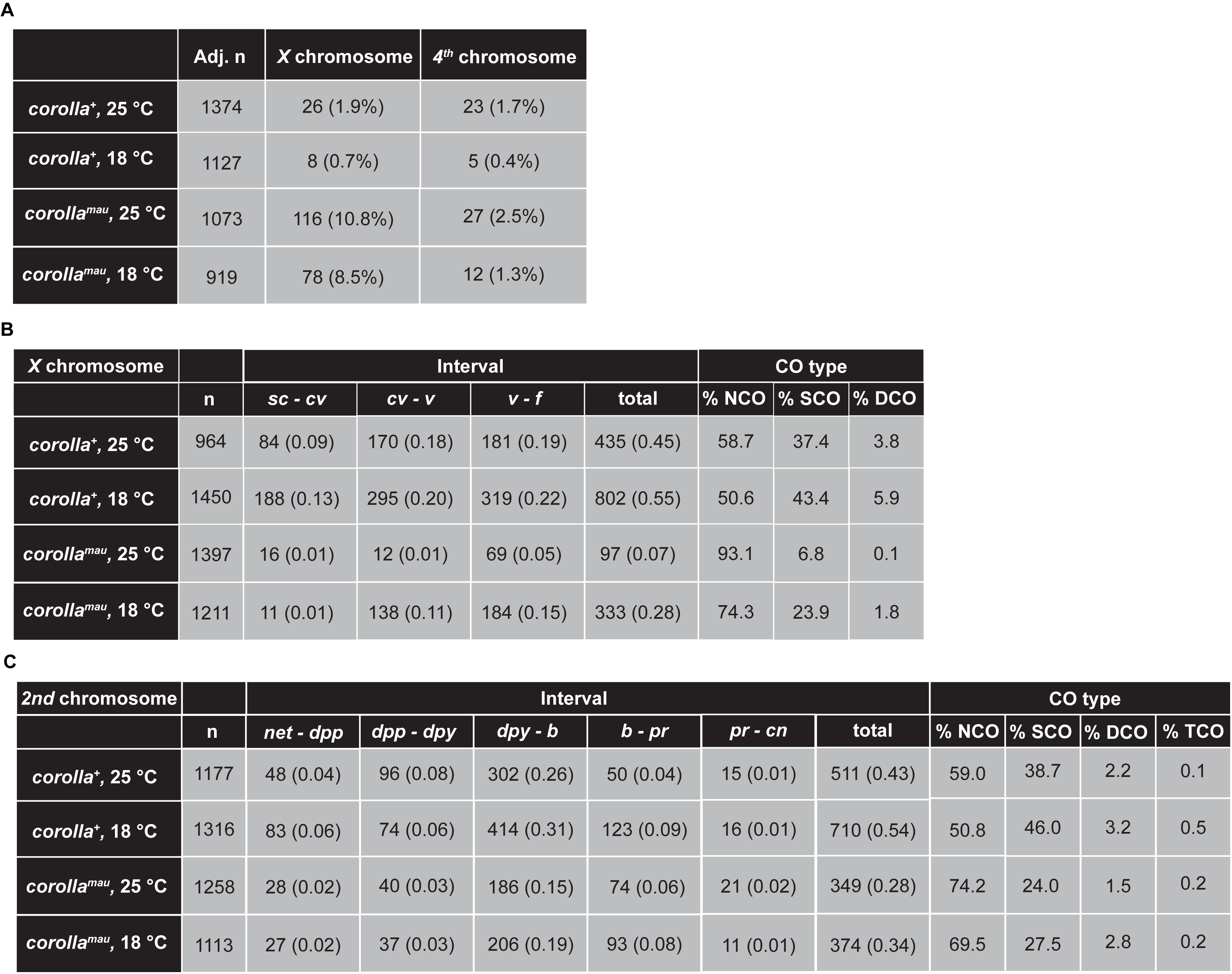

### Supplemental Figure 9

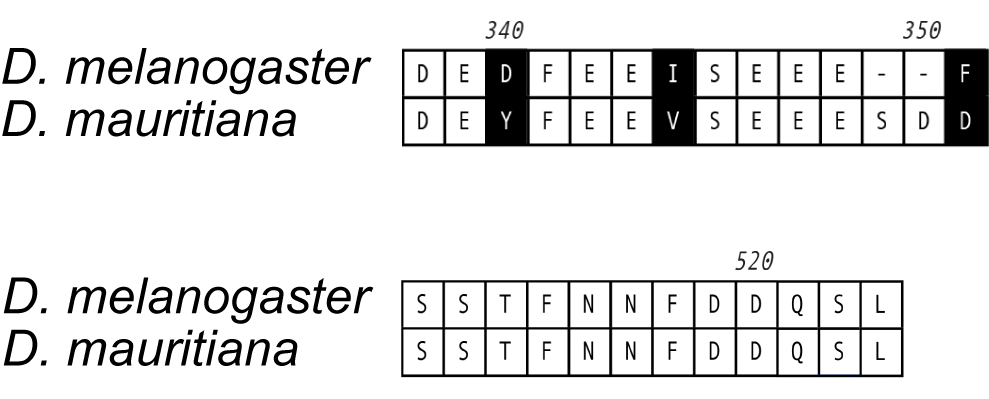

### Supplemental Figure 10

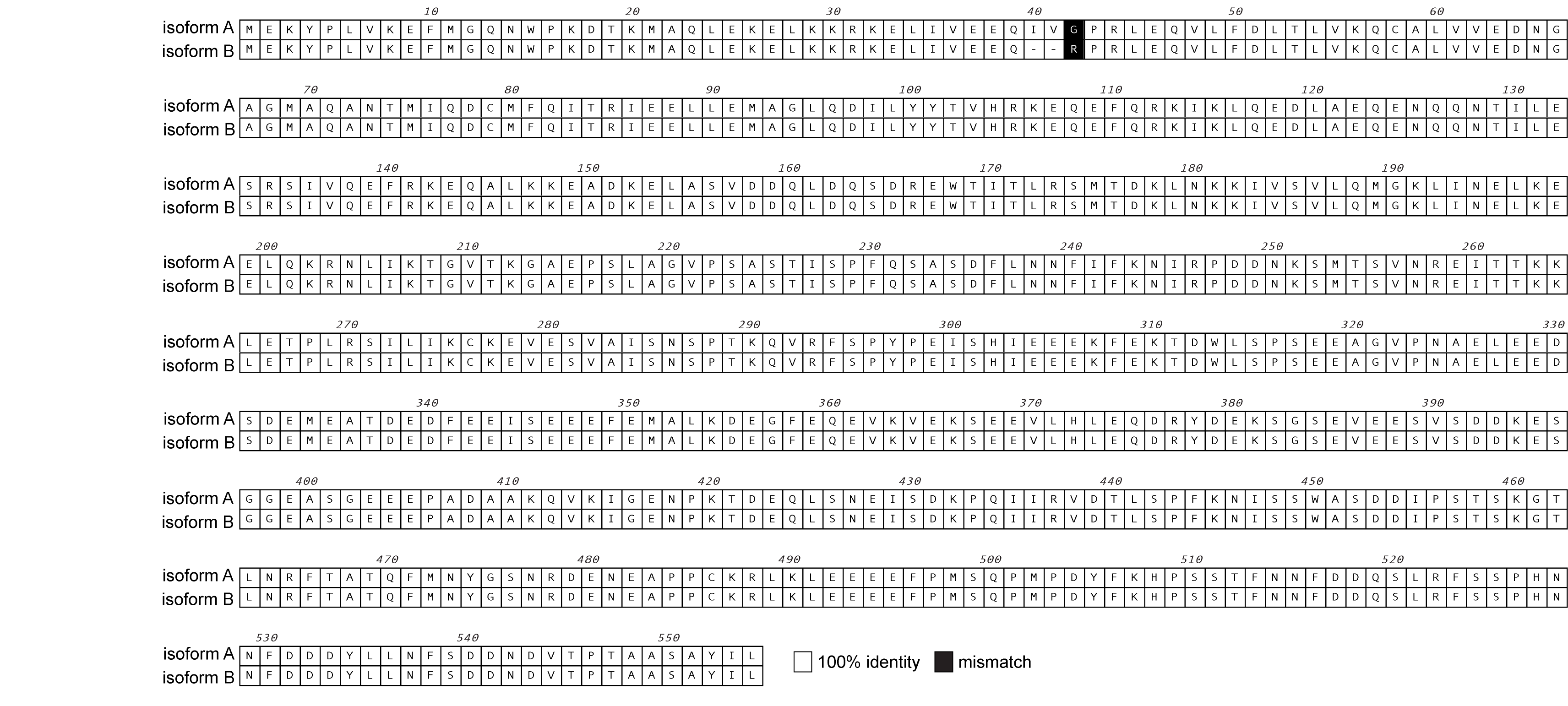

### Supplemental Figure 11

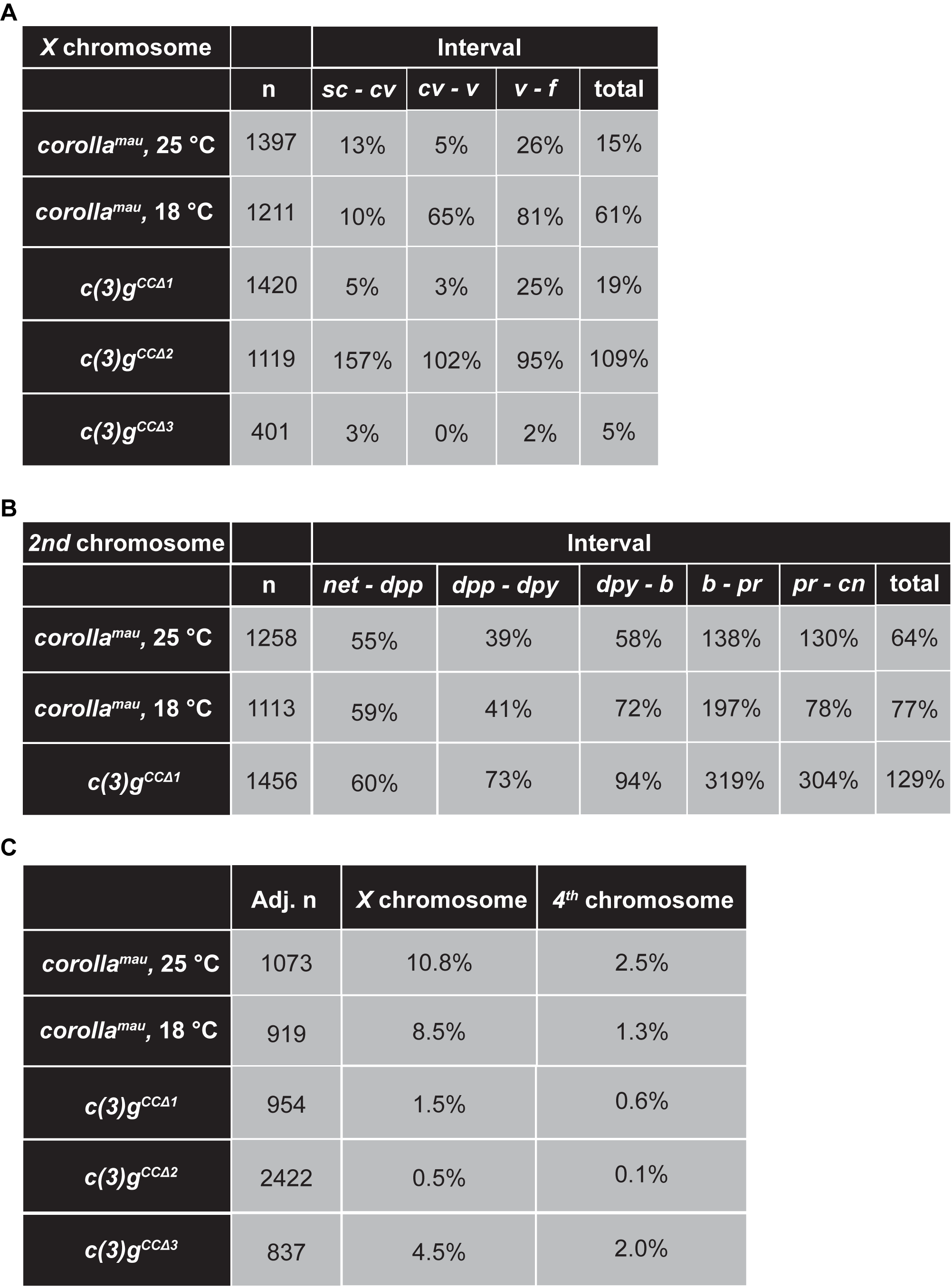
