## Supplemental File 1 for "Using evolution as a tool: Replacing *corolla* in *Drosophila melanogaster* with its *Drosophila mauritiana* ortholog creates a novel hypomorphic allele": S1_file copy.rtf

ClustalW multiple sequence alignment

2 Sequences Aligned           Processing time: 0.7 seconds
Gaps Inserted = 1             Conserved Identities = 482
Score = 4756

Pairwise Alignment Mode: Fast
Pairwise Alignment Parameters:
    ktup = 1   Gap Penalty = 3   Top Diagonals = 5   Window Size = 5
    Similarity Matrix: blosum

Multiple Alignment Parameters:
    Open Gap Penalty = 10.0   Extend Gap Penalty = 0.2
    Delay Divergent = 30%     Gap Distance = 4
    Similarity Matrix: blosum

1. D. melanogaster vs. D. mauritiana

   Aligned Length = 556   Gapped Segments = 1   Total Mismatches (including gaps) = 74
   Identities = 482 (86.7%)   Similarities = 26 (4.7%)

D. melanogaster   1 MEKYPLVKEFMGQNWPKDTKMAQLEKELKKRKELIVEEQIVGPRLEQVLFDLTLVKQCAL  60
D. mauritiana     1 MEKYPLVTEFMGQNWLKDTQMAQFENELRKRKELIVEEQIIGPQLEQVLFDLTLAKQCAL  60

D. melanogaster  61 VVEDNGAGMAQANTMIQDCMFQITRIEELLEMAGLQDILYYTVHRKEQEFQRKIKLQEDL 120
D. mauritiana    61 AVENNGAGLAQVNTMVQDCIFQITRIEELLEMAGLQDILYYTMHRKEQEFLHKIKLQEDL 120

D. melanogaster 121 AEQENQQNTILESRSIVQEFRKEQALKKEADKELASVDDQLDQSDREWTITLRSMTDKLN 180
D. mauritiana   121 AEQENQQNTILESRSIVQELRKEQVSKKEADKELASVDDQLDQADREWTITLRSMTDNRN 180

D. melanogaster 181 KKIVSVLQMGKLINELKEELQKRNLIKTGVTKGAEPSLAGVPSASTISPFQSASDFLNNF 240
D. mauritiana   181 KKIVSVLQMRKLINELKEELQKRNLIKTVVTKGAEPHLAGVLSAGSISSFQSASDFLNNF 240

D. melanogaster 241 IFKNIRPDDNKSMTSVNREITTKKLETPLRSILIKCKEVESVAISNSPTKQVRFSPYPEI 300
D. mauritiana   241 IFKNIKLDGNKSMTSVNSEITTKKLETPLRSILVKCKEDEPVAISNSPTKQVRFSPYPEI 300

D. melanogaster 301 SHIEEEKFEKTDWLSPSEEAGVPNAELEEDSDEMEATDEDFEEISEEE--FEMALKDEGF 358
D. mauritiana   301 SHIEEDKFEKPDWLSRSEEADVPNAELEEDADEIEATDEYFEEVSEEESDDEMELKDEGF 360

D. melanogaster 359 EQEVKVEKSEEVLHLEQDRYDEKSGSEVEESVSDDKESGGEASGEEEPADAAKQVKIGEN 418
D. mauritiana   361 EQEENVEKSEKLLDPEQDRYDDKSGSEVEESVSDDKESGGEASGEEEPADAAKQVKFGEK 420

D. melanogaster 419 PKTDEQLSNEISDKPQIIRVDTLSPFKNISSWASDDIPSTSKGTLNRFTATQFMNYGSNR 478
D. mauritiana   421 PQPDEQPSNEISDKPQIIRVDTLSPIKNISSWTSDDIPTTSKGTLNRFTAAQLMNYGSDR 480

D. melanogaster 479 DENEAPPCKRLKLEEEEFPMSQPMPDYFKHPSSTFNNFDDQSLRFSSPHNNFDDDYLLNF 538
D. mauritiana   481 DANEEPPSKSLKLEEDEFPMTQPMPDYFKHPSSTFNNFDDQSLRFSSPHNNFDDDYLLNF 540

D. melanogaster 539 SDDNDVTPTAASAYIL 554
D. mauritiana   541 SDDNDVTPTAASAYIL 556
